# Roost switching and behavioural shifts following human disturbance of vampire bats in complex landscapes

**DOI:** 10.1101/2025.05.20.655023

**Authors:** Rita Ribeiro, Carlos M. Zariquiey, Wendi Chavez, Kristhie Pillaca Rodriguez, Joaquin Clavijo Manuttupa, Marco Antonio Risco Huiman, Roselvira Zuniga Villafuerte, Walter Campana Quintanilla, Noelia Medrano-Uchuya, Hollie French, Jakub Czyzewski, Carlos Shiva, Nestor Falcon, William Valderrama, Daniel P. Walsh, Tonie E. Rocke, Jason Matthiopoulos, Daniel G. Streicker

## Abstract

In Latin America, management of vampire bats has had inconsistent effects on the spread of lethal rabies infections from bats to humans and livestock. Positive and negative effects on rabies are hypothesised to reflect changes in bat roost fidelity and behaviour following human disturbance, but high landscape complexity has precluded monitoring bat movement in the areas where rabies transmission and bat management activity are most intense. We used animal-borne GPS tags to track 93 vampire bats in a rabies hotspot in the Peruvian Andes, where limited satellite visibility led to partial observation of animal locations. Immediately after capture, handling and tag application, 43% of bats, particularly males and bats with longer forearms, disappeared from the study area, suggesting a rapid roost switching response. Among the remaining 55 bats, a Bayesian state-space model revealed high individual heterogeneity on the disturbance night but on average increased flight distances. Over the remaining 8-night study period, bats gradually remained closer to the roost. These disturbance-driven shifts provide mechanisms to explain how bat disturbance can both accelerate and decelerate rabies spatial spread. More broadly, we present a modelling framework to infer key movement metrics in landscapes where GPS data limitations challenge conventional movement models.

## Introduction

The composition and geomorphological structure of landscapes determine how animal hosts move within their environment and therefore how infectious diseases spread. For example, large urban areas shape the movement of bobcats (*Lynx rufus*) and the transmission of their Feline Immunodeficiency Virus (Retroviridae: Lentivirus), while in less urban areas, the same virus in puma (*Puma concolor*) is structured by mountain ranges and other landscape features that determine host-relatedness and spatial connectivity ^1,2^. Pathogen spread within or across complex landscapes can directly influence management decisions. For instance, the fungal pathogen *Batrachochytrium dendrobatidis* has spread through rugged mountainous landscapes in Latin America, causing major amphibian declines. Hence, the capacity to forecast its spatial and temporal spread would greatly improve monitoring and prevention efforts ^3^. However, human activities intended for disease control and/or management of reservoir hosts may themselves also alter animal behaviour in ways that are counterproductive to management. For example, culling of badgers (*Meles meles*) for control of bovine tuberculosis (*Mycobacterium bovis*) altered the ranging and dispersal behaviours of surviving badgers, increasing tuberculosis risk in cattle outside of culled areas ^4^.

In Latin America, rabies transmitted by vampire bats (*Desmodus rotundus*) illustrates how uncertainty in animal movement across complex landscapes can hinder the control of an important disease. In this region, vampire bats are the main reservoir of rabies virus (Rhabdoviridae, Lyssavirus), which causes sporadic lethal outbreaks in humans and a sustained agricultural burden through livestock mortality ^5^. In addition to vaccination campaigns targeting humans and livestock, rabies management throughout the region includes sanctioned use of anticoagulant poisons (‘vampiricides’) to reduce bat population sizes and unsanctioned killing of captured bats and destruction, sealing or burning of roosts ^6–8^. However, because long term persistence of vampire bat rabies depends on bat movement, it has been hypothesised that dispersal of surviving bats might enhance rather than reduce rabies spread ^6^. Indeed, evaluation of a culling campaign in Peru showed that culling vampire bats in response to livestock rabies outbreaks increased viral spatial spread, whereas culls carried out before rabies arrival appeared to slow its spread ^7^. Both effects were hypothesised to arise from increased bat movement, but the behavioural responses of vampire bats to direct human disturbance remain speculative.

Directly measuring fine-scale behaviour in vampire bats is impeded by technical and ecological challenges. In mark-recapture studies, low between-roost recapture rates limit inference on movement ^9^. Very high frequency (VHF)-radio tracking has been applied, but is labour-intensive, error-prone and only records animal presence near antennas ^10–12^. The recent adoption of smaller, lighter and longer-lasting animal-borne Global Positioning System (GPS) tags has revolutionised our understanding of animal movements that may underlie pathogen spread ^13^. In principle, GPS tracking of vampire bats could test how landscape geomorphology and direct human disturbance influence bat movement, connect colonies and enable rabies virus spread into new areas.

However, the reliance of GPS tags on satellite acquisition remains a major limitation in the rugged mountainous or densely forested landscapes where vampire bats occur and where rabies transmission is most problematic ^14,15^. To our knowledge, no GPS study of vampire bat movement has been reported.

Here, we used lightweight GPS tags with remote data recovery to quantify how vampire bats respond to direct human disturbance (i.e., capture, handling, and tagging). We focused on the Apurimac, Ayacucho and Cusco (AAC) departments in the Peruvian Andes, which is composed of subtropical valleys (>1200-3600 meters) that are inhabited by bats, separated by precipitous peaks exceeding 6000 meters ^14^. Although the mechanism is not understood, this landscape complexity likely contributes to AAC being the primary hotspot of vampire bat rabies in Peru, accounting for 60.2% (annual range, 32.9-81.4%) of all reported livestock cases between 2003 and 2019, despite regular bat culling campaigns ^7^. However, the same landscape complexity that appears to promote rabies circulation also reduces satellite acquisition by GPS tags, creating a major hurdle for examining how human disturbance alters bat movement in a key context for rabies management. To address this challenge, we developed a hierarchical Bayesian state-space model that combines a movement process model with an observation-error model, allowing for the explicit propagation of uncertainty due to missing fixes and location error ^16^. We jointly estimated two movement metrics, nightly departure times and distances from the roost, while accounting for GPS error. We used this model to identify the ecological drivers of departure times and maximum flight distances from roosts, focusing on the hypothesis that the disturbance of capture, handling and tagging would induce longer flights. We further investigated the persistent disappearance of many bats from the focal roosts immediately after capture, handling and GPS tagging.

## Methods

### Ethics and permits statement

The study, including bat capture, handling and tagging, received ethical approval from the University of Glasgow Research Ethics Committee (Reference EA28a/21) and research permits from Peruvian authorities, including SERFOR (Servicio Nacional Forestal y de Fauna Silvestre, Nos. D000024-2022, D000107-2022, D000103-2024 and D000188-2024-MIDAGRI-SERFOR-DGGSPFFS-DGSPFS) and SERNANP (Servicio Nacional de Áreas Naturales Protegidas por el Estado, N° 021-2023-SERNANP-SHM/J). All procedures followed ARRIVE guidelines.

### Study area and design

The study was conducted during the Peruvian dry season, between April-October 2022 and May-June 2023, in five vampire bat roosts (four caves and one abandoned house, Supplementary Table S1), each located >50 km apart in southern Peru (Fig. 1a). At each roost, two tagging sessions were carried out, separated by at least three months. Roost 2 (Fig. 1c) had a single tagging session due to data recovery challenges. Each session aimed to tag 10 adult bats (five males, five females). All capture sessions began after dusk, except the first capture in April 2022, when, due to logistical constraints, captures began in the morning.

**Figure 1.**
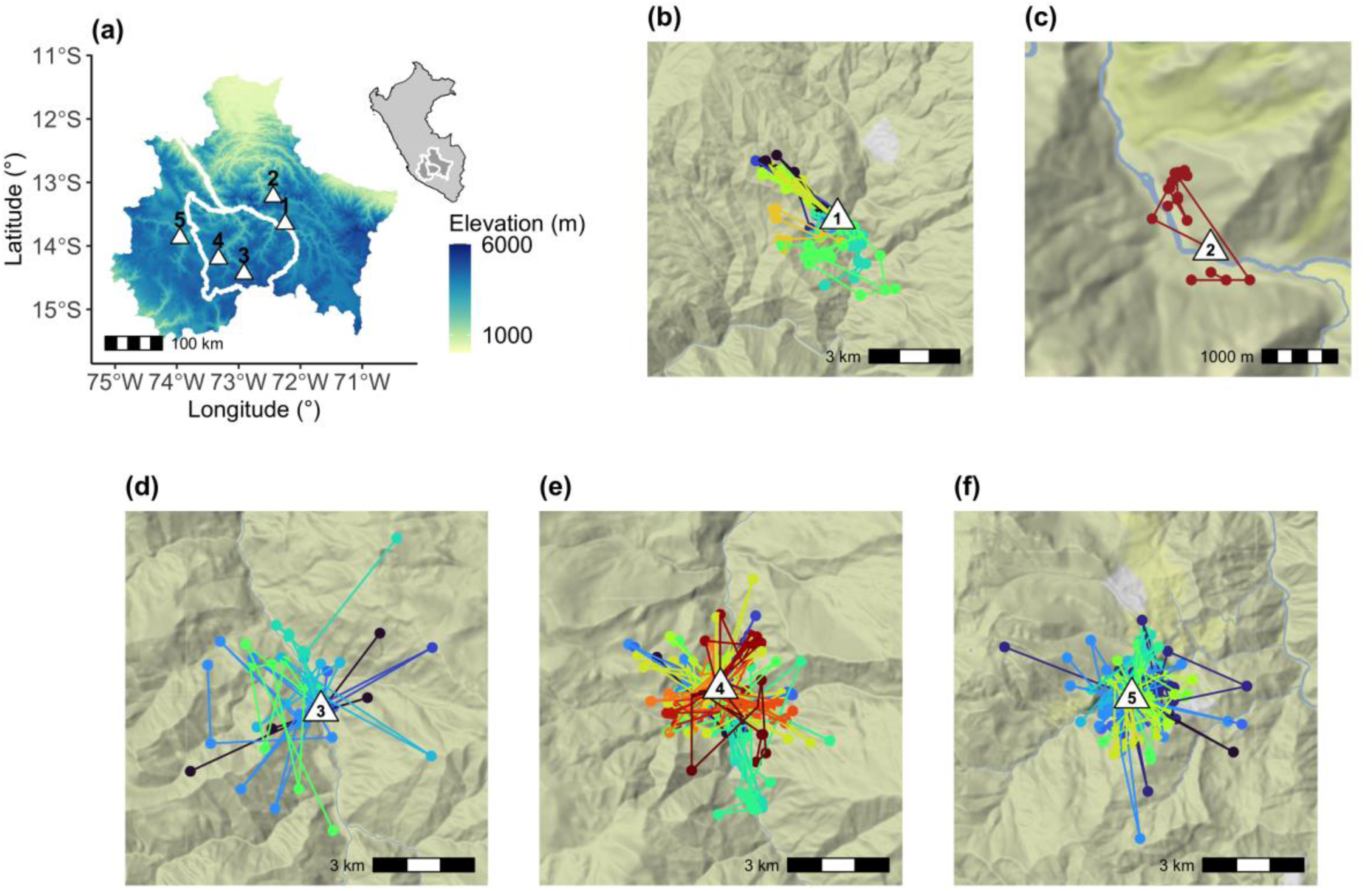
Study area and GPS data from 55 vampire bats (*Desmodus rotundus*) from five roosts. (a) Elevation map with the five study roosts (numbered triangles) in AAC, with an inset map of Peru highlighting the study region. (b) roost 1, (c) roost 2, (d) roost 3, (e) roost 4 and (f) roost 5. In panels b-f, each colour represents a bat tracked for up to nine nights. Due to the varying scales of movement across the roosts, this figure does not use a standardised scale, and only 95% of the fixes closest to the roost were plotted to avoid outliers. Maps used ggmap ^55^. Base map tiles show terrain; in panel (c) the map is zoomed in. Data © OpenStreetMap contributors, © Stadia Maps, © Stamen Design, © OpenMapTiles. Licensed under CC BY 3.0 / ODbL (https://www.openstreetmap.org/copyright).

During initial deployment in 2022, GPS tags were set to record locations every 5 minutes (10 tags) or 10 minutes (five tags) for three and six nights, respectively. To extend tracking to nine nights in subsequent sessions, later deployments used a lower sampling frequency of 15 minutes. All tags were tested at fixed locations in open environments, and some were additionally evaluated in different environments of the study area using both stationary and mobile trials. All tags were fully charged before deployment. Tags operated nightly from 6 PM to 6 AM, with data downloaded remotely via one or two solar-powered UHF receivers (PathTrack, UK) placed at the roost entrance. GPS fix (‘position’) success is often diminished in enclosed environments where wild animals rest ^17^. In each deployment, one or two tags were intentionally left inside the roost, confirming that tags did not acquire fixes when bats remained inside the roosts.

Short-term follow-up recapture sessions (one to two nights) were conducted at the end of the GPS-tracking study to determine whether bats with no recovered GPS data had left the roost or were present but no longer carrying a functioning tag. Data from capture sessions carried out 4-32 months later for ongoing monitoring projects (hereafter, ‘long-term’ recapture sessions), were used to compare long-term recapture rates of bats that disappeared versus returned to the roost after disturbance (Supplementary Table S1). In all the sessions, roosts were inspected for dead bats and lost tags.

### Capture, handling and tagging of vampire bats

Vampire bats were captured using mist and butterfly nets. Captured bats were fitted with a 3.5 mm wing ring (Porzana, UK) for individual identification, and weight (g), sex, reproductive status (non-reproductive, scrotal male, pregnant, or lactating female), age (adult, subadult, or juvenile), and forearm length (mm) were recorded. GPS tags (2.8 g, 3 cm body plus 5 cm antenna; nanoFix®, PathTrack, UK) were applied to bats that met the inclusion criteria (see below). After light sedation using intramuscular (pectoral) injection (29-gauge needle, 1.0 mL syringe) of Ketamine (20–25 mg/kg) diluted in saline (NaCl) at a 50/50 volume ratio or Midazolam (0.25 mg/kg, used in 2023 to allow faster recovery times), GPS tags were attached to the skin of the dorsal side of each bat using an absorbable suture (Monosyn®, B Braun), threaded through custom tag loops ^18^. According to the manufacturer, the absorbable sutures lose 50% of their strength after 14 days, allowing natural tag detachment. Before suturing, hair was trimmed, and a small amount of cyanoacrylate-based glue was used to secure the tags. Sedation and tag suturing were performed by a veterinarian, ensuring it did not impair flight or walking. GPS tags were painted with UV fluorescent paint to aid tag recovery (Starglow®, Glowtec, UK).

Only healthy adult bats were GPS-tagged, while pregnant and lactating females were excluded for animal-welfare reasons. The average weight of tagged bats was 43.4 g (± 3.4 g, N=93), 44.1 g (± 3.3 g, N=43) for females and 42.7 g (± 3.4 g, N=50) for males. For both sexes, the tags represented a maximum of 7% of the bat’s weight (mean = 6.4 ± 0.47% for females and mean = 6.6 ± 0.47% for males). Recommendations for bat tracking studies indicate that attached devices should not exceed 10% of body mass, with a best practice of limiting to 5% ^19^. However, heavier tags may be acceptable for short-term use in species with high physiological tolerance for weight change, as seen in bats ^20–23^. Our previous records of wild adult vampire bat mass (N=3,429 individuals) showed that they can gain over 9 g of weight after feeding (pre-feeding: 41.9 g ± 8.44, N=2,751; post-feeding: 51.5 g ± 9.1, N=678), implying daily weight gains of >20% of body mass. Additionally, females can exceed 85 g during pregnancy and even forage while carrying pups. These weight fluctuations and their ability to carry significant additional mass strongly indicate that the tags we used should be tolerable for short-term use.

### GPS data cleaning and management

GPS fix quality increases with the number of satellites acquired: one or two satellites produce very inaccurate 1-dimensional (1D) fixes, three satellites produce inaccurate 2-D fixes, and four or more satellites produce accurate 3-D or 3-D+ fixes ^15,17,24^. According to the manufacturer, nanoFix GPS fix success and precision depend on satellite availability, geometry, and signal quality, and a minimum of four satellites is required to generate a location; otherwise, no fix is generated. Consequently, all positions obtained are 3-D or 3D+ fixes and represent the highest-quality fixes available from the device.

GPS data were retrieved for 60 of the 93 tagged bats. Across all bat-nights (12 hours, from 6 PM to 6 AM), the overall raw fix success (percentage of attempts which retrieved a location) was 11%. However, this estimate included bats whose GPS tags returned only a few fixes before the animals disappeared from the study area, as well as periods when bats were inside roosts, where fixes were not generated (see above). To obtain a more biologically meaningful estimate, we restricted the calculation to bats retained for subsequent modelling (see below) and quantified the proportion of successful fixes during the bats’ active period, defined as the interval between the first and the last recorded fix for each bat-night. Within this proxy of active period, we observed an average fix success of 56% (Supplementary Fig. S1), which is within the range of reported for other mammal GPS studies (43-99% ^15,17^).

As part of data processing, any GPS fixes recorded prior and during tag application (within 1-hour post-capture) were removed. Data from the first night were otherwise included to assess immediate disturbance behaviour. Additional filtering removed one inaccurate fix flagged by the software, data from one malfunction tag, individuals with insufficient data (< 3 fixes), one unrealistically stationary tag (i.e., suggesting tag loss or bat death) and unrealistic flight speeds ^25^. Full details of data cleaning are provided in supplementary information (S2). To manage the three sampling rates (5, 10 and 15 minutes) and provide a uniform input to the Bayesian state-space model, each GPS fix was assigned to a 15-minute block during the night (i.e., 48 15-minute blocks). Data were processed in R version 4.4.0 ^26^.

### Drivers of post-disturbance roost switching

The return of GPS-tagged bats to their original roosts was monitored using UHF receivers placed at roost sites and through recapture sessions conducted immediately after the tracking period (i.e., short-term sessions). For bats with no GPS detections from the receivers and no recaptures during the short-term sessions, the most likely explanation is that they did not return to the roost during the nine-night study period. This pattern is consistent with roost switching, although alternative explanations (e.g., detection failure, tag loss, mortality) cannot be fully excluded.

We fitted binomial generalised linear models (GLMs) to examine whether the probability of returning to the roost after the night of disturbance was random or associated with individual bat traits (i.e., sex and forearm length). Because vampire bat weight fluctuates considerably within a night depending on feeding and time since feeding, bat weight was not included. Nevertheless, we included the percentage of body mass represented by the tag to evaluate the potential influence of tag burden on roost switching behaviour. Roost identity was included as a fixed effect to account for baseline differences among roosts. We assessed multicollinearity using variance inflation factors (threshold of 2.5 ^27^) and compared model performance across candidate models using the Akaike information criterion (AIC) and pseudo-R^2^. The significance of added terms and interactions was evaluated using nested analysis of variance.

### Variable definition and processing for the GPS analysis

The state-space model investigated predictors of bats’ nightly departure time and distance travelled from the roost. Vampire bats are thought to be lunar phobic, departing before moonrise or after moonset ^28^. Therefore, departure time was modelled as a function of the number of hours until complete darkness, which depended on the interplay of moonrise, moonset, astronomical night start (occurs after sunset and dusk) and end (occurs before dawn). For each roost and night, the hours from sunset until darkness were calculated using the suncalc package (v0.5.1) ^29^. We predicted that longer periods until darkness would delay departure times. We also tested whether sex influences departure time.

Distance from the roost was modelled as a function of variables related to roost location, individual bat characteristics, and human disturbance. Roost-related variables within 5 km radii of roosts included proportion of permanent water, predicted vampire bat roost density ^30^ and cattle density ^31^ (Supplementary Table S2), calculated using the packages sf (v1.0-19) and terra (v1.7-71) ^32,33^. We predicted that the availability of alternative roosts and preferred food sources (cattle) near the roost would reduce the distances travelled. Since cattle graze near water and vampire bats can use rivers as corridors, we hypothesised that permanent water sources near roosts could aid prey searching, decreasing distances from the roost ^34^. Individual characteristics included bat sex and forearm length. We hypothesised that males would fly further distances after direct disturbance due to reduced offspring care ^35^. We also expected that individuals with longer forearms may travel farther from the roost or would be more likely to switch roosts after capture, since forearm length scales with wingspan and area, providing a proxy for wing loading and flight speed ^36,37^, and is positively associated with migration distance in bats ^38^.

Human disturbance was defined as the effects of capture, handling, and tagging procedures. Key variables included the night of capture (’night 0’) versus subsequent nights, and the time post-disturbance, calculated as the difference in hours between the start of the tracking session and each GPS fix date and time stamp. Time post-disturbance for each bat and each night was averaged to obtain a single value per bat per night. Averaging reduced the influence of irregular GPS fix intervals and temporal autocorrelation, and provided a covariate structure consistent with our model, which included effects of individual, roost and night. We anticipated that direct disturbance would increase flight distance following capture, with distances decreasing over time as behaviour normalised. Finally, the number of rural settlements within 5 km of each roost was used as a proxy for historical disturbance levels (e.g., culling, roost destruction or shifts in agricultural practices), allowing us to isolate the effects of the immediate disturbance caused by our capture, handling and tagging from long-term historical persecution effects (Supplementary Table S2).

### Bayesian state-space model

A Bayesian state-space model was fitted to the vampire bat GPS data with three objectives: (1) to estimate latent movement metrics, including times and distances from the roost; (2) to evaluate how bat traits, environmental variables, and direct human disturbance influenced these metrics; and (3) to smooth partially observed GPS tracks in order to obtain robust estimates of the latent states under measurement error.

The Bayesian state-space model included two latent variables: the bats’ departure time (*t*, in minutes since the daily activation of the GPS at 6 PM) and its distance from the roost (ψ, in km). The model inferred distances from the roost every 15 minutes and estimated the duration of nightly activity and return time to the roost, accounting for both process error (i.e., the natural stochasticity in the biological processes) and observation error (i.e., GPS measurement error). To estimate the latent states (i.e., times and distances), the state-space model requires a fixed reference point, which we assumed to be the known roost. Although vampire bats may use multiple roosts ^39^, our analysis focuses on bats that returned to the focal roost. A preliminary analysis of the GPS data supported this assumption, indicating that at least 57-64% of bat-nights included a return to the focal roost (Supplementary Fig. S2). We further note that these estimates are likely conservative as rapid returns or signal loss at the roost can prevent some fixes from being recorded. We also assumed that most bats would depart before 1 AM, as 95% of our experiments were conducted on nights when the moon set before 1 AM.

For each bat *i* and night *j, t*_*i,j*_ is the departure time in minutes after 6 PM, with a maximum value of 420 minutes (1 AM). The expected and realised departure instances, τ^-^ _*i,j*_ and τ^-^_*i,j*_, are expressed as a proportion of the interval between 6 PM and 1 AM,

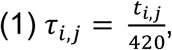

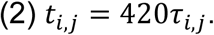

The expected departure instance τ^-^ _*i,j*_ was modelled as a logit function of the number of hours until darkness and bat sex,

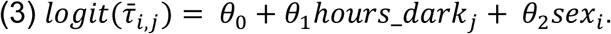

τ^-^_*i,j*_ was modelled using a Beta distribution with mean τ^-^ _*i,j*_ and a process error quantified as the standard deviation of departure instances σ. This error represents variability in trip start times and was assigned a wide prior (estimated by the model, Supplementary Table S3), to capture pooled inter-trip and inter-individual variation,

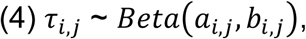

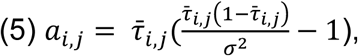

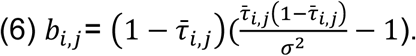

The duration of nightly activity after departure was modelled with a uniform distribution, assuming that once a bat departed, it remained away for at least 20 minutes. The maximum duration of a trip was the time left after departing and the end of GPS activation (720 minutes),

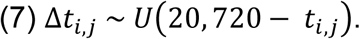

To estimate distances travelled from the roost, we defined a parametric model of distance against time, for each bat *i*, night *j*, and each block *k* of 15 minutes after 6 PM,

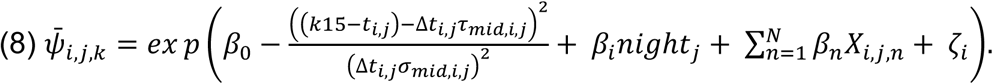

This equation assumes that only one trip is made by each bat every night (i.e., the bat leaves the roost, travels to a destination and then comes back), but is flexible regarding specific details of the trip, such as when it starts, ends, and maximum distance from the roost. It also assumes that the net velocity is away from the roost until the maximum distance is reached, and then toward the roost thereafter, for 15-minute intervals. The exponential of the intercept β_0_, corresponds to the maximum distance from the roost. β_*i*_ corresponds to the terms for the effect of night of disturbance (*night*_*j*_) for each bat, and β_*n*_ is the parameter for the n^th^ covariate effect (forearm length, sex, roost density, number of human rural settlements, cattle density, permanent water and time post-disturbance). We added an individual-level random effect ζ_*i*_ with mean 0 and standard deviation σ_1_ (ζ_*i*_ = *N*(0, σ_1_)), to capture the influence of individual variation (Supplementary Table S3). In both departure time and distance models, we considered random effects of individual, roost, calendar night, and session of capture. Only the random effect of the individual on the distance model was relevant, hence it was kept.

The mid-point of the trip τ^-^_*mid,i,j*_ was defined as the proportion of time between departure and return until reaching maximum distance, with a parameter σ_*mid,i,j*_, which made sure that the distance from the roost climbed smoothly from the start of the trip to the midpoint and then declined to the trip’s endpoint, preventing unrealistic jumps in movement. τ^-^_*mid,i,j*_ was given a beta prior distribution and was set to be close to the middle of the trip duration (mean 0.5 and variance 0.001). σ_*mid,i,j*_ was fixed to a sufficiently large value to ensure that the bats did not immediately reach their maximum distance after departure or instantaneously returned to the roost,

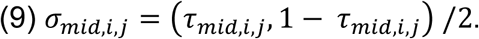

The realisation of distance giving rise to observations, ψ_*i,j,k*_, was modelled using a normal distribution, with mean ψ^-^_*i,j,k*_ and variance *O*_*i,j,k*_,

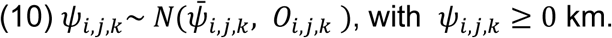

*O*_*i,j,k*_ is a combined error that incorporates observation error ξ_*i,j,k*_, the standard deviation of GPS error, and process error, in the form of an overdispersion parameter ф, to account for further uncertainty that cannot be explained by the observation model (i.e., in equation 11, 0.6 km is the GPS error for four satellites, the minimum number of visible satellites required),

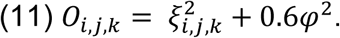

The overdispersion parameter ф was estimated by the model, but we encouraged it to be close to zero (Supplementary Table S3) to prioritise the identification of biological effects. The nanoFix manufacturer indicated using the number of visible satellites to estimate GPS error. Using data from a GPS tag placed at a single location in our study area, we fitted a regression of the standard deviation of distances against the number of satellites and estimated the standard deviation of GPS error for each satellite count (standard deviation of 0.3 km when four satellites were visible; deviance explained = 0.986). This regression produced the two parameters γ_0_ and γ_1_ which, in combination with the number of visible satellites that generated each positional fix, *N*_*i,j,k*_, informed ξ_*i,j,k*_

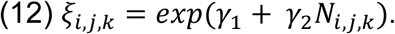

For variable selection, we used Bayesian Stochastic Search Variable Selection (SSVS) with spike and slab priors to estimate model coefficients conditional on a variable being retained in the model ^40^. If a variable was participating in the model, the coefficient value was given a normal distribution, with mean 0 and standard deviation τ^-^_1_, with τ^-^_1_ being estimated by the model, except for the coefficient for hours until darkness, which was given a gamma prior distribution to preserve its positivity (mean 0.01 and variance 0.0001). The posterior for the probability of retention ρ was modelled with a Bernoulli distribution (variable included or excluded), with the parameter ρ being estimated by the model with a constrained Beta prior (i.e., a variable has a 25% probability of being included), meaning that variables need strong evidence to be included.

We assessed goodness of fit by calculating the R^2^, comparing observed distances from the roost to the posterior predicted distance, both for the entire model and for each bat separately. The models were fitted in JAGS 4.3.2 using rjags (v4-15) and runjags (v2.2.2-4) ^41,42^. The model ran with four independent chains, for 10,000 samples per chain after discarding the first 500,000 iterations as burn-in, with thinning of 200 (retaining every 200^th^ iteration). Convergence across chains was assessed through visual inspections of the posterior and the Gelman-Rubin diagnostic test (using coda (v0.19-4.1) ^43^). Supplementary Table S3 lists the final model parameter priors and posteriors.

## Results

Between April 2022 and June 2023, 93 adult vampire bats were GPS-tagged across five roosts in southern Peru. Three bats were confirmed dead and at the end of the study, GPS data was obtained for 60 bats (60/90, 67% of tagged bats). No GPS data was recovered from 30 bats, and nine additional bats were only detected (i.e., with GPS fixes) during ‘night 0’, indicating the possibility that 43% (39/90) of bats left the focal roosts following direct disturbance and did not return during the 9-night tracking period. The disappearance rate from night 0 (43%) was higher than that observed in subsequent nights, suggesting an acute response to disturbance rather than typical roost switching behaviour (Supplementary Fig. S3). If the apparent disappearance of bats reflected GPS tag loss or failure, we would expect missing bats to have similar short-term recapture rates as bats that returned. However, of the 39 bats which disappeared on the disturbance night, we recaptured only one (short-term recapture rate = 2.6%) during the short-term re-capture sessions (one to two nights immediately after the GPS tracking session). In contrast, of the 51 bats that remained in the study area (as evidenced by GPS data upload to receivers stationed at roosts), eight were recaptured during the short-term sessions (short-term recapture rate = 15.7%). In contrast, recapture rates during the long-term sessions (4-32 months after GPS tagging) were similar between groups (missing bats: 15.4% [N=6] versus GPS detected bats: 11.8% [N=6], arguing against bat mortality as a driver of disappearance during the GPS tracking. We therefore interpret the disappearance of bats as most compatible with a failure to return to the roost over the duration of the GPS tracking session (i.e. roost switching).

A binomial GLM showed that bats with longer forearms (−1.10 [Standard error (SE) = 0.42], P<0.01) and male bats (−1.55 [SE = 0.83], P=0.06) were less likely to return to the roost after the direct disturbance, although there was only weak evidence for the latter effect. Disappearance occurred in all roosts and differed significantly among them. Due to this roost-level variation, we checked whether it influenced the individual-level effects; however, the effects of forearm length and sex were unchanged when roost was removed, indicating they were not driven by roost-level differences (Fig. 2; Supplementary Table S4; Supplementary Fig. S4). Across all models tested, including those with interaction terms, we found no evidence that the tag burden influenced bat disappearance (Supplementary Table S5).

**Figure 2.**
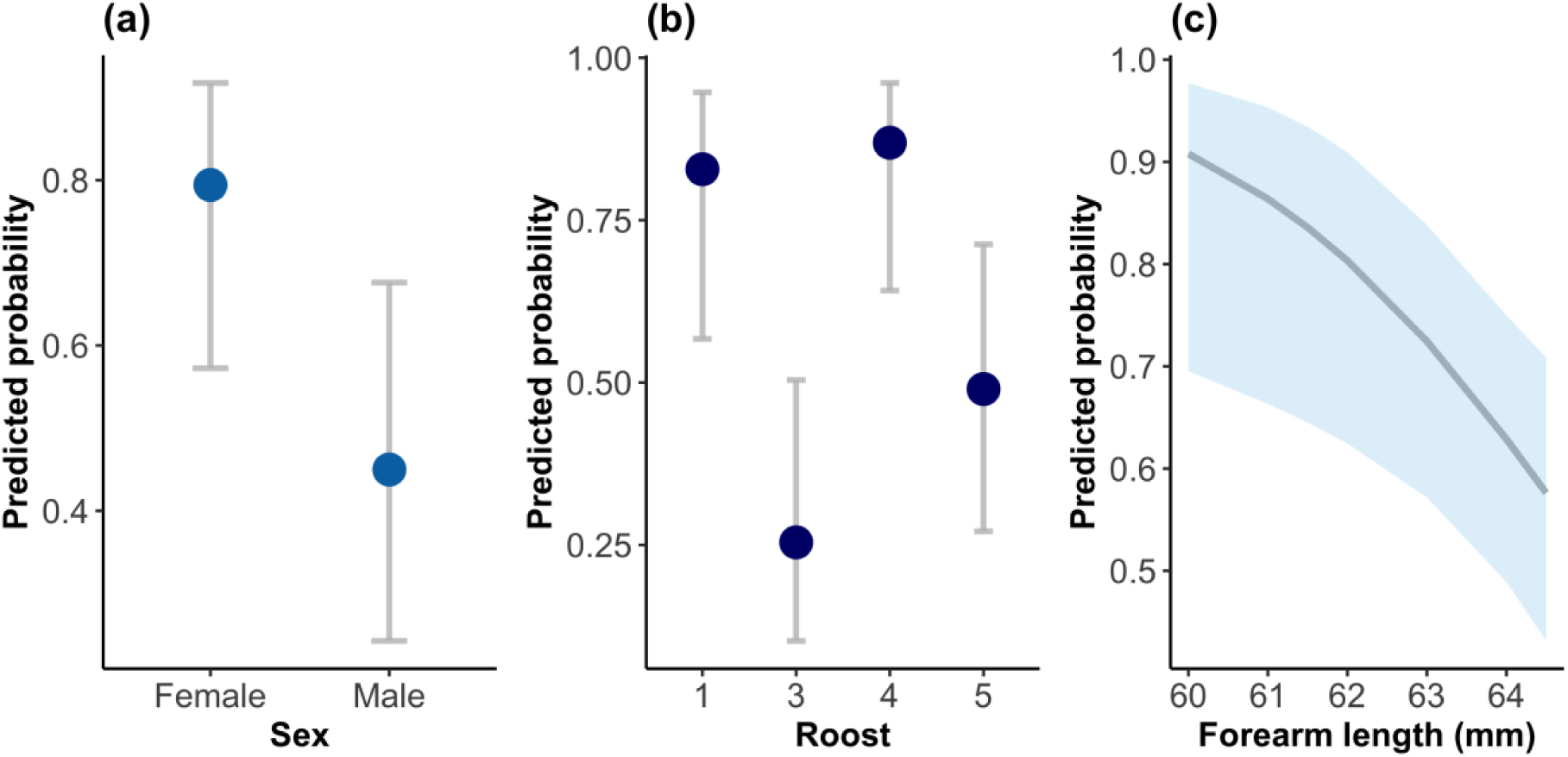
Predicted probabilities of vampire bats (*Desmodus rotundus*) returning to the roost following disturbance. Results from a binomial GLM: (a) Shows sex-specific estimates with 95% confidence intervals; (b) shows the roost-specific estimates with 95% confidence intervals (roost 2 was not included in this analysis) (c) shows the relationship of probability of return with forearm length (mm), with the 95% confidence interval shaded.

Visualisation of the GPS data from individuals that returned close enough to roosts to upload GPS data showed that vampire bats occasionally flew to distances > 5 km from the roost but spent most time < 5 km from roosts (Fig. 1b-f; Supplementary Fig. S5-S6). Individual bats from the same roost foraged in different areas, did not visit the same locations every night, and used a variety of flight paths, highlighting behavioural heterogeneity among and within individuals (Fig. 1b, d, e, f; Supplementary Fig. S7).

We next explored correlates of departure time and maximum nightly distance from the roost in 55 bats with GPS data that met our inclusion criteria (see data cleaning above and in S2). Each individual contributed between one to nine nights of data, yielding a total of 242 bat-nights. Our main hypothesis was that direct human disturbance would lead to longer flights. The Bayesian state-space model demonstrated a strong overall fit to the GPS-derived distance data (R^2^ = 0.80, Supplementary Fig. S8a). Because the number of observed GPS fixes varied across bats, we also calculated individual-level R^2^ values to assess model fit consistency. The model fitted well for most bats (median individual R^2^ = 0.71), and only 10.9% of distance profiles showing poor fit (R^2^ < 0.20) (Supplementary Fig. S8b-S12). Although predicted maximum distances exceeded observed maximum distances per bat-night, this arises because the model estimates latent distances during periods without observations (Fig. 3c-d).

**Figure 3.**
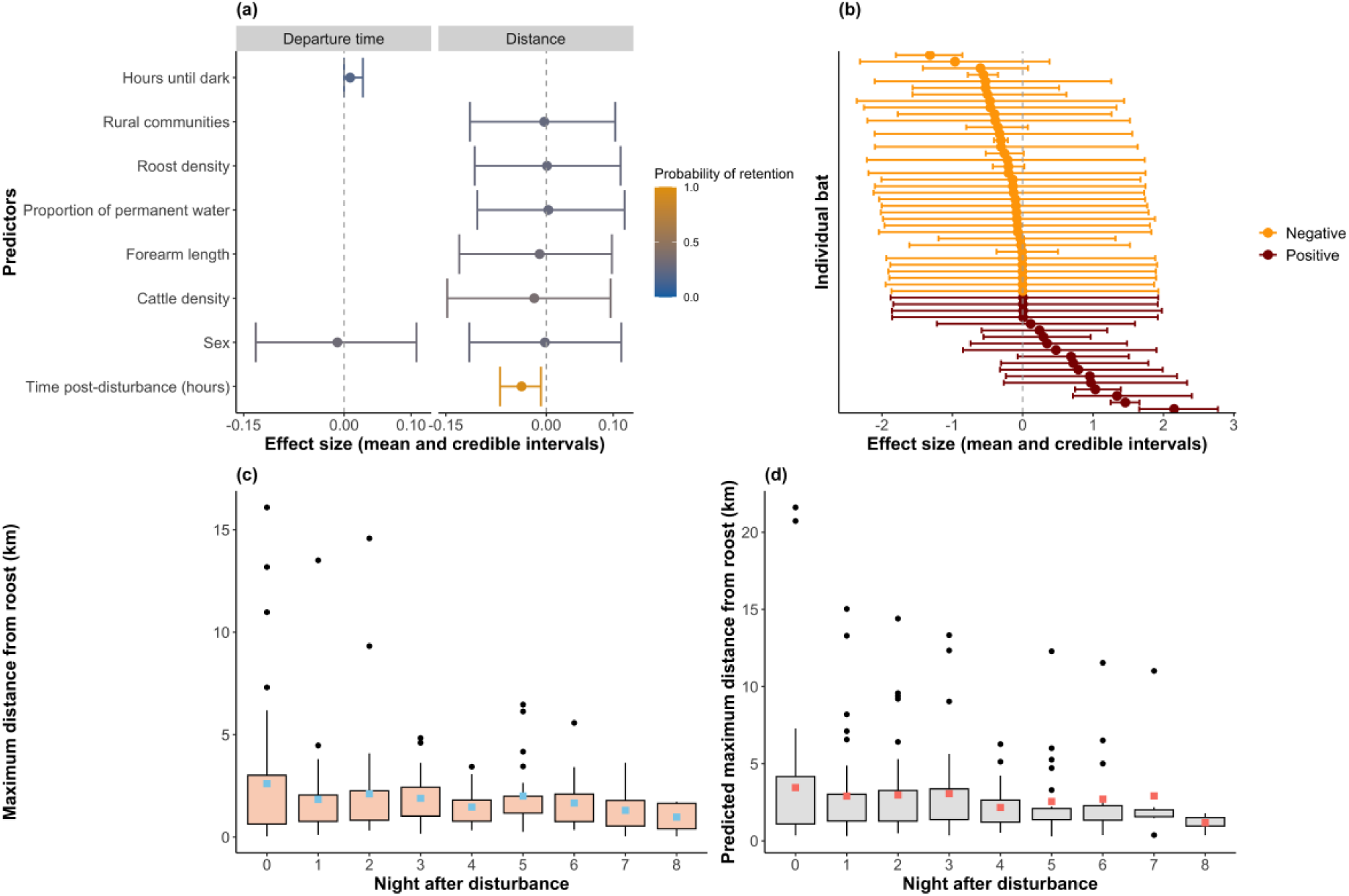
Predictors of vampire bat (*Desmodus rotundus*) departure time and distance from the roost. (a) Standardised effect sizes on the log-odds scale (departure time, logit model) or log scale (distance, exponential model), coloured by SSVS retention probability. (b) Individual variation in the distance from the roost on the disturbance night (bat-night interaction). (c) Observed maximum distances (km) from the roost per bat and night. (d) Predicted maximum distances (km) per bat and night. In (c) and (d), night 0 is the disturbance night; squares indicate means, and black dots are outliers.

Our analysis of departure times showed that, on average, bats departed ∼2 hours after the GPS started at 6 PM (posterior mean of 124 minutes, 95% credible interval (CI) [58.3; 220.5]) and were out of the roost for >5 hours (posterior mean of 345 minutes, 95% CI [190.0; 497.4]) (Supplementary Fig. S9). Although the number of hours until darkness had a low posterior inclusion probability, its posterior distribution indicated that bats left the roost as soon as there was no sunlight or moonlight (Fig. 3a; Supplementary Table S6).

Among drivers of the nightly distances that bats moved from the roost, time post-disturbance was the only variable with strong posterior support for inclusion (probability = 0.95, Fig. 3a, d; Supplementary Table S6). This negative effect implied that bats initially flew farther from the roost following disturbance, then shifted to flying shorter distances, presumably returning to a more typical flight distance. Specifically, the maximum nightly distance decreased by 0.17 km per night (Supplementary Fig. S10-S11) and stabilised approximately four nights after tagging (Fig. 3d). Deeper examination of model outputs also showed an interaction between individual bat identity and night (night of disturbance versus all the other nights), whereby on the night of the disturbance, some bats stayed unusually close to the roost, and others undertook the longest movements recorded in the study period (Fig. 3b-4; Supplementary Table S7; Supplementary Fig. S12-S13). This putative individual heterogeneity in the response to disturbance occurred across all roosts except roost 2, which had data from only one bat (Supplementary Fig. S14). Taken together, these results suggest that while on average bats flew further from the roost on the night of disturbance, this result was dampened by marked individual heterogeneity, where some bats had the opposite response of reducing movement.

**Figure 4.**
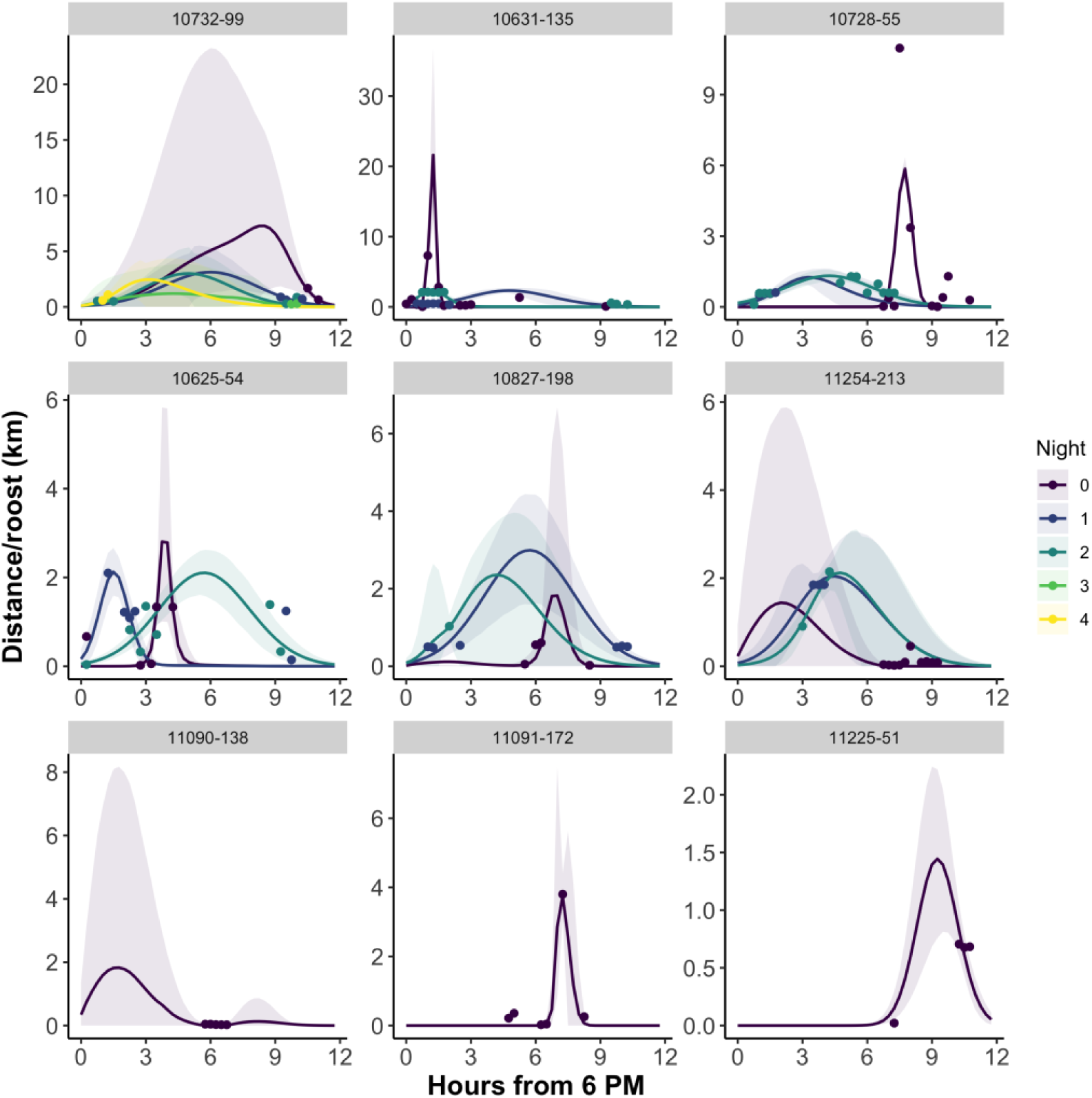
Reconstructed distance-over-time curves for nine vampire bats (*Desmodus rotundus*) across nine nights. First row: bats that travelled longer distances on night 0; middle row: bats that travelled similar distances across different nights; bottom row: bats that left the roost after night 0 and did not return. The shaded area indicates the credible intervals, and the dots the observed values. Purple lines indicate night zero, with progressively lighter colours indicating later nights. Due to the varying scales of movement, the y-axis is not standardised.

## Discussion

Patterns of rabies maintenance and spillover from vampire bats are thought to be strongly linked to bat dispersal, yet vampire bat movement has been notoriously difficult to observe. By GPS tracking 93 vampire bats in a rabies hotspot in the Peruvian Andes, we revealed new insights into the natural history of this key viral reservoir in Latin America. We also identified three behavioural responses of vampire bats to direct human disturbance: elevated roost switching, increased nightly flight distances, and high individual heterogeneity which could influence rabies transmission and the effectiveness of bat culling as a control strategy.

High-resolution GPS tracking of vampire bats revealed patterns that were previously unobservable in mark-recapture or radiotelemetry studies. Previous work has suggested that vampire bats primarily move along valleys and rivers, following downstream corridors shaped by landscape ^39,44^. However, our GPS data show that, rather than consistently foraging downstream, vampire bats also crossed mountainous terrain and foraged in both upstream and downstream directions.

Furthermore, similar to other colonial species such as seabirds, vampire bats exhibited high movement behavioural heterogeneity within the colony ^45^. Earlier work suggested that vampire bats repeatedly used the same foraging sites and prey, yet our GPS data reveal much more variable nightly movements, challenging this assumption ^46^. Overall, heterogeneity across individual bats and within bats over short periods helps explain the sometimes-sporadic evidence of vampire bat bites on livestock and humans ^47^ and implies that pathogen-infected bats may expose multiple animals at different locations. Incorporating these behavioural ecological insights on vampire bats into models of rabies spread could enhance our understanding of its dynamics.

Following disturbance 43% of the GPS-tagged bats apparently disappeared from the monitored study area for the duration of our tracking study, a pattern that could not be explained by tag failure or mortality. Disappearance was further elevated for males (which may have greater flexibility to switch roosts due to lack of offspring care) and bats with longer forearms (which may have greater dispersal capacity) further supports roost switching as the most plausible explanation for disappearance (Fig. 2) ^35–38,48^. As vampire bats are known to use multiple nearby roosts within their home range, a key uncertainty from our analysis is whether the rate of disappearance we observed reflects normal or elevated roost switching behaviour ^39,49^. Although we could not include an undisturbed control roost because measuring returns to roosts required GPS tagging, the magnitude of disappearances observed immediately after disturbance compared to that observed in subsequent nights of GPS tracking supports an acute disturbance effect over normal behaviour (Supplementary Fig. S3). Future work using less invasive methods, such as radio frequency identification (RFID)-based automated systems, could help establish baseline roost-switching rates and quantify the magnitude of the disturbance effect suggested by our data.

Among the 55 GPS-tagged bats that remained detectable, comprising a total of 242 bat-nights, we observed longer distance movements following direct disturbance, and higher individual heterogeneity on the night of capture. As with the roost switching analysis, the need for GPS tagging meant that baseline flight distances in the absence of disturbance could not be inferred. Although we interpret the decline on flight distances over time to reflect a gradual return to typical behaviour, it is alternatively conceivable that data from the initial nights after capture and tagging represented normal behaviour and the shorter distances observed at the end of the study period were influenced by tag effects such as fatigue. However, we suggest that high individual heterogeneity on the night of disturbance (Fig. 3b) and the pattern of roost switching that we observed (Fig. 2, Supplementary Fig. S3), are more consistent with an acute disturbance response ^50^. Furthermore, previous work has shown that terrestrial mammals can exhibit acute behavioural changes immediately after capture and tagging, with some species increasing their movement distances, although the magnitude and direction of this effect vary across species and contexts ^51^. Bats were not included in these analyses, and our results extend this work by revealing marked individual-level variation within a single population exposed to the same disturbance event.

As a pest species and zoonotic reservoir, vampire bats experience intense persecution through sanctioned population control campaigns using anticoagulant poisons, alongside unsanctioned control via informal methods and roost destruction. Collectively, our results show these disturbances likely induce acute changes in vampire bat behaviour which might alter the spread of rabies virus in complex and context dependent ways. For example, in areas where rabies is already circulating, disturbance-induced roost switching or increased movement could increase rabies transmission by increasing overlap in foraging areas and mixing between colonies, thereby facilitating the spread to nearby roosts. In contrast, large-scale disappearance of bats from key roosts in rabies-free populations could reduce the likelihood of viral establishment by creating more fragmented populations composed of smaller, more isolate colonies. Since male bats appeared more likely to disperse following disturbance, these effects may be especially relevant given that viral spatial spread is believed to be strongly influenced by male bat movement ^48^. Consistent with these hypotheses, a phylogenetic analysis of rabies spread in the context of vampire bat culling showed that reactive culls (i.e., in rabies infected areas) accelerated spread, while preventive culls (i.e., in rabies free areas) delayed arrival of the virus ^6,7^. Our results therefore reinforce the risks of widely practiced reactive and/or unsanctioned management of rabies outbreaks and highlight the need for improved models to anticipate risk to empower preventive action. As vampire bat population control is expected to continue, we encourage coordinated efforts across health sectors and affected communities and the development of reproductive suppressants as an alternative to culling which may both promote herd immunity and reduce behavioural responses ^52^.

GPS fix loss in terrestrial systems is often non-random, with up to 37% of fixes missing due to predictable habitat-driven constraints rather than random noise. Dense canopy, complex terrain, or enclosed structures, reduce satellite acquisition, and these effects may be further modulated by animal behaviour, producing additional variability ^15,17,24^. These predictable environmental effects, and their interaction with bat behaviour (i.e., small mammals that fly and roost in caves and other enclosed structures), explain much of the data sparsity in our study. Because habitat driven effects are systematic and measurable, it justifies the selection of a state-space model for estimating distances travelled accounting for observation error and uncertainty from missing fixes ^15,16^. The Bayesian framework we developed jointly estimated movement parameters across 55 GPS-tracked bats. Although GPS data were partially observed, the dataset comprised bats from five roosts and 242 bat-nights, a relatively large sample for bat GPS studies, which typically involve few individuals and short tracking periods ^21–23^. Robust ecological inference from GPS studies depends not only on the number of fixes but also on the number of animals sampled ^53^, making this dataset strong despite partial data. By partially pooling information across bats, the model leveraged well-sampled individuals to inform estimates for those with sparser data, while propagating uncertainty arising from missing observations ^16^. Importantly, our objective was not to reconstruct fine-scale movement trajectories from sparse data, but to infer how individual, environmental, and disturbance-related factors influence coarser key movement metrics that remain identifiable under such sampling, particularly maximum distances travelled from the focal roost (Fig. 3c-d). These ‘true distances’ that animals can travel, specially under different circumstances, such as human disturbance, are critical for disease management, but are often missed in sparse datasets, which tend to underrepresent maximum distances ^54^.

A potential limitation of our modelling approach is the assumption that vampire bats make a single nightly trip and return to the focal roost. Although 13% of bat-nights showed bimodal posterior distributions consistent with multiple trips per night (Supplementary Fig. S12), the strong correspondence between observed and predicted distances (Fig. 4, Supplementary Fig. S8-S12) and convergence diagnostics (Supplementary Table S3-S6-S7) support the single-trip approximation as a reasonable simplification. A second assumption is the use of a single roost as the reference point, even though vampire bats may use multiple roosts ^39,49^. This reflects a model requirement for a fixed origin and is conditional on bats returning during the tracking period. Return was confirmed for 57-64% of bat-nights (Supplementary Fig. S2), though this may be an underestimate due to data sparsity. Model fit and convergence further indicate that the focal roost served as an effective reference point for most movements. If some long-distance movements reflected unobserved roost switching, our results could overestimate true foraging distances. However, because data recovery required eventual return to the focal roost, these movements still represent maximum displacements. Biologically, such distances remain relevant regardless of mechanism, as they still create opportunities for livestock contact and rabies transmission.

Human disturbance can reshape wildlife movement and pathogen spread, yet its effects remain poorly understood in geomorphologically complex landscapes, where terrain both influences transmission and challenge the study of disease reservoirs. Our study shows how advances in GPS technology, combined with appropriate analytical frameworks, can extend wildlife tracking to species and environments where conventional analyses may fail, or be strongly biased by satellite visibility constraints. Moreover, we demonstrate that direct human disturbance can have repercussions for vampire bat rabies management, revealing movement dynamics that would otherwise remain unknown in complex landscapes and highlighting how altered bat dispersal and colony mixing may facilitate the emergence of livestock rabies in new areas of Latin America.

## Acknowledgements

We are grateful to Elsa Cardenas Canales, Haris Malik and Carlos Tello for field assistance; Lynda Dunbar and Marieke Pingen for help with the suturing technique; Fred Touzalin and Pathtrack for insights and assistance with the GPS. We thank Holly Niven for helpful discussions about the analysis, and Nardus Mollentze, Travis Mcdevitt-Galles, and our colleagues from the Streicker group for peer review and feedback on earlier versions of the manuscript. We thank SENASA (National and Regional offices) for feedback and support.

## Author contributions

Conceptualisation: R.R., J.M., D.G.S. Methodology: R.R., J.C., J.M., D.G.S. Analysis and visualisation: R.R., J.M. Investigation: R.R., C.M.Z., W.C., K.P.R., J.C.M., M.A.R.H., R.Z.V., W.C.Q., N.M-U., H.F., D.G.S. Resources and management: C.M.Z., C.S., N.F., W.V. Data Curation: R.R. Project administration: D.G.S., Funding acquisition: D.P.W., T.E.R., J.M., D.G.S. Supervision: R.R., J.M., D.G.S. Writing (original draft): R.R., J.M., D.G.S. All authors contributed critically to the drafts and gave final approval for publication.

## Data availability statement

Data and code for this study are deposited in the Dryad Digital Repository and will be available after manuscript publication.

## Consent for publication

Not applicable.

