## Supplementary for "Roost switching and behavioural shifts following human disturbance of vampire bats in complex landscapes"

[5] Escuela Profesional de Ciencias Biológicas. Facultad de Ciencias, Universidad Nacional de Piura, Piura, Perú

[6] Centro de Investigación Vertebrate, Universidad Nacional San Antonio Abad del Cusco, Cusco, Perú

[7] Medical Research Council-University of Glasgow Centre for Virus Research, Glasgow, UK

[8] School of Biodiversity, One Health and Veterinary Medicine, BioElectronics Unit, University of Glasgow, Glasgow, UK

[9] Facultad de Medicina Veterinaria y Zootecnia, Universidad Peruana Cayetano Heredia, Lima, Perú

[10] U.S. Geological Survey, Montana Cooperative Wildlife Research Unit, Wildlife Biology Program, University of Montana, Missoula, Montana, USA

[11] U.S. Geological Survey, National Wildlife Health Centre, Madison, Wisconsin, USA

### Supplementary Information

**Contents:**

**Table S1:** Summary of vampire bat (*Desmodus rotundus*) captures for the GPS study.

**Figure S1:** Fix success per individual bat during a proxy of the active period.

**S2:** Supporting information for data cleaning and filtering.

**Table S2:** Variables used in the Bayesian state-space model.

**Figure S2:** Nightly return status across 242 bat-nights.

**Table S3:** Model parameter priors and hyperpriors, and posteriors.

**Figure S3:** Bat disappearance after the direct disturbance (nights 0-1) and bat detection loss across all the other nights.

**Table S4:** Results for the binomial generalised linear model on drivers of vampire bat (*Desmodus rotundus*) roost switching.

**Figure S4:** Assessment of performance of binomial generalised linear model M12.

**Table S5:** Results for the binomial generalised linear model on drivers of vampire bat (*Desmodus rotundus*) roost switching - interactions with bat traits.

**Figure S5:** Spatial distribution of vampire bat (*Desmodus rotundus*) GPS locations for five roosts in southern Peru.

**Figure S6:** Distribution of distance from the roost across roosts.

**Figure S7:** Individual GPS data for vampire bats (*Desmodus rotundus*) for up to nine nights of tracking.

**Figure S8:** Model goodness of fit for correlates of bat departure time and nightly distance from the roost.

**Figure S9:** Predicted times from the Bayesian state-space model.

**Table S6:** Summary of the Bayesian state-space model for vampire bats (*Desmodus rotundus*).

**Figure S10:** Changes in the maximum predicted travel distance of vampire bats (*Desmodus rotundus*) over nights post-disturbance.

**Figure S11:** Changes in the maximum predicted travel distance of vampire bats (*Desmodus rotundus*) from the roost.

**Table S7:** Posterior summaries of the interaction coefficients between 55 vampire bats (*Desmodus rotundus*) and night of disturbance.

**Figure S12:** Reconstructed distance-over-time curves for 55 vampire bats (*Desmodus rotundus*).

**Figure S13:** Nightly individual distance effects.

**Figure S14:** Coefficient of variation of observed distance of vampire bats (*Desmodus rotundus*) from the roost by roost.

**Table S1: Summary of vampire bat (*Desmodus rotundus*) captures for the GPS study.**

| Roost | [Latitude, Longitude] | Region | Session | Date | Number of bats captured | Number of tagged bats | Tagged males | Tagged females | Dates of short-term recaptures | Dates of long-term recaptures[1] |
|---|---|---|---|---|---|---|---|---|---|---|
| **1** | *[-13.65257, -72.24618]* | *Cusco* | *1* | *20/10/22* | *17* | *10* | *3* | *7* | 30/10/22 | 12-13-14/03/24 |
| | | | 2 | *10/06/23* | *15* | *8* | *6* | *2* | 20/06/23 | |
| **2** | *[-13.2241762, -72.4402573]* | *Cusco* | *1* | *13-16/10/22*[2] | *25* | *9* | *4* | *5* | 26/10/22 | 14-15/06/23; 07-08-09/03/24 |
| **3** | *[-14.439, -72.91237]* | *Apurimac* | *1* | *15/08/22* | *37* | *10* | *5* | *5* | 24/08/22 | 16-18-19/03/24; 28-29-30/06/24; 08-09-12-14/10/24; 30/04-01-02/05/2025 |
| | | | 2 | *14/05/23* | *40* | *10* | *3* | *7* | 23-24/05/23 | |
| **4** | *[-14.203246, -73.32493]* | *Apurimac* | *1* | *25/04/22* | *26* | *15* | *11* | *4* | 05-09/05/22 | 18/05/23; 02-03-04/03/24 |
| | | | 2 | *21/08/22* | *30* | *10* | *5* | *5* | 02/09/22 | |
| **5** | *[-13.8772911, -73.9500989]* | *Ayacucho* | *1* | *23/09/22* | *22* | *10* | *4* | *6* | 03/10/22 | 05-06-10/11/23; 23-24-25/03/24; 04/07/24; 18-19-20/10/24; 22-23-26/04/25 |
| | | | 2 | *08-12-21-24/06/23*[3] | *24* | *11* | *9* | *2* | 18-19/06/23 | |

[1]Long-term captures for all Session 1 events (except at Roost2) include the Session 2 captures and recaptures. Roosts with captures expanding into 2025 (Roosts 3 and 5) include RFID antennas, which necessitate long-term monitoring.
[2]This capture session was split across two nights because heavy rain on the 13th limited capture success. To meet the target of 10 bats per session, additional captures were performed on the 16th.
[3]Low capture success at the cave in 2023, required supplementary captures to reach the target sample size. Recaptures were limited by logistical constraints at the end of fieldwork.

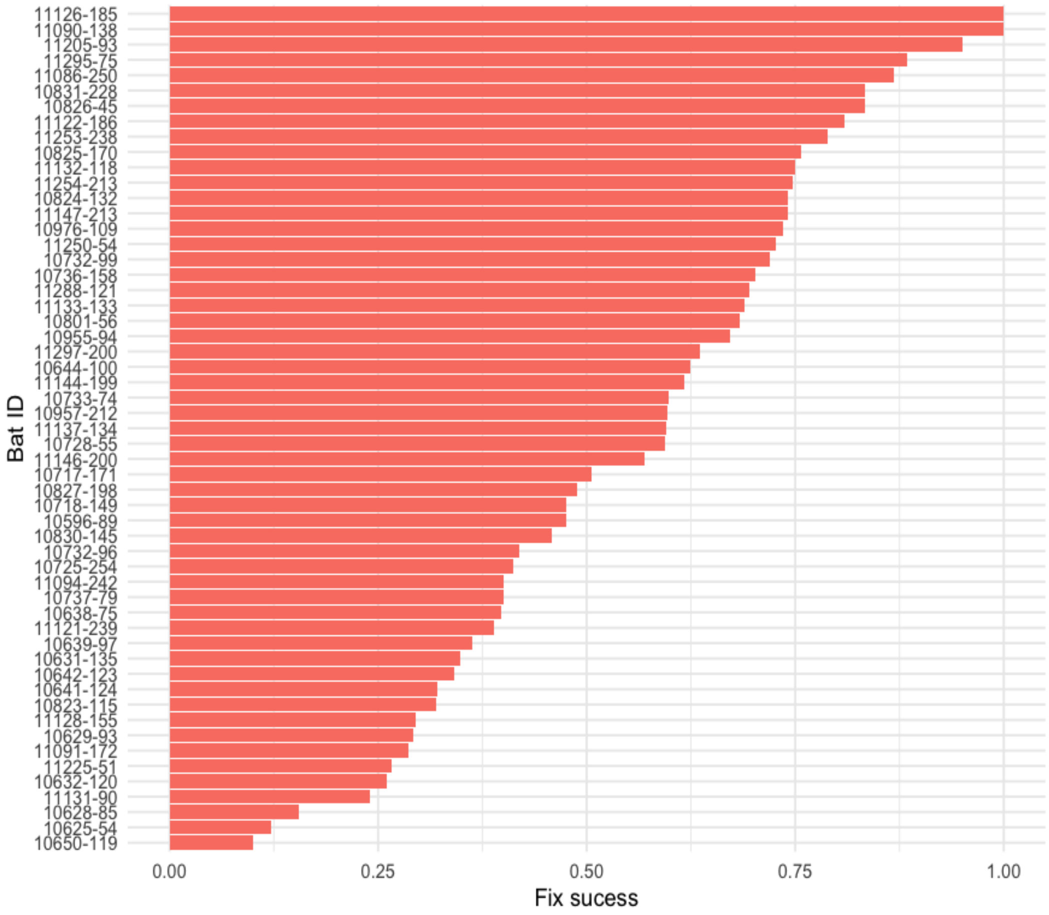


**Figure S1: Fix success per individual bat during a proxy of the active period.**

Among the 60 bats with recovered GPS data, 89% of possible observations (14,142 out of 15,842) were not recorded due to having less than four visible satellites (i.e., 11% fix success), leaving a total of 1,700 GPS fixes. To obtain a more biologically meaningful estimate, we restricted the calculation to bats retained for subsequent modelling and quantified the proportion of possible fixes that were successful during the bats' active period, defined as the interval between the first and the last recorded fix for each bat-night. Within this proxy of active period, we observed an average fix success of 56% (Fig. S1).

**S2: Supporting information for data cleaning and filtering.**

GPS data filtering included one inaccurate fix flagged by the software, all fixes from one bat with a malfunctioning tag (distances up to 6 km recorded before attachment), all data from three bats with less than three fixes across the study period, all data from an unrealistically sessile bat (suggesting tag loss or death), GPS fix jumps and outliers of estimated flight speed [1]. The speed (km/hour) of each segment between subsequent fixes was calculated, along with the Euclidean distance from the roost for each fix, using base R and the sf package (v1.0-19) [2].

Maximum flight speeds for vampire bats are unclear. Sánchez-Hernández *et al.* [3] reported average speeds of 13.82km/h (range 9.6 - 27.3km/h) for males and 13.36km/h (range 7.2 - 23.4 km/h) for females under artificial conditions of capturing and releasing individual bats inside a tunnel. In contrast, Daler *et al.* [4] suggested maximum speeds of up to 20 m/s (72 km/hour) based on a robot modelled after *D. rotundus*. We removed two instances where two subsequent segments had a high-speed value (i.e., ~ 20 km/hour), with the bat returning to the same position at the same speed, indicating GPS error rather than actual movement. We also removed two fixes (GPS locations) associated with two speed outliers, estimated at 72.8 and 115.7 km/hour, respectively. The remaining fixes included speeds up to 57.6 km/hour; however, close inspection of the data suggested that the bats were engaged in active travel rather than returning directly to the same position after reaching a high speed. Our state-space model was designed to be robust to GPS errors, so we could have omitted data filtering based on speed entirely. However, we anticipated that simultaneously inferring and filtering speed within the model could hinder convergence.

**Table S2: Variables used in the Bayesian state-space model.**

Variables (units, spatial resolution and source) used to estimate bat departure time, activity duration, and travel distances from roosts in southern Peru from April 2022 to June 2023. Variables were grouped into three categories: individual, roost or disturbance related. NA means 'not applicable'.

| Variable (units) | | Spatial resolution | Source |
|---|---|---|---|
| Individual | Forearm length (mm) | NA | Field records |
| | Sex (1 for females and 0 for males) | NA | Field records |
| Disturbance | Night of disturbance (1) versus all the other nights (0) | NA | Field records |
| | Time post-disturbance (hours) | NA | Field records |
| | Number of rural human settlements (count) | 2 $km^2$ | Obtained from Ribeiro et al., [5]. Original data from Peru Government (INEI and CEPLAN). |
| Roost related | Roost density (count/ $km^2$) | 8 $km^2$ | Spatial predictions of vampire bat roost density from Ribeiro et al., [5]. |
| | Proportion permanent water | 10 $m^2$ | Landcover 2021 [6]. |
| | Cattle density (count/ $km^2$) | ~ 10 $km^2$ | Rescaled for ~ 5 $km^2$. Global distributions of cattle. FAO data, 2015 [7]. |

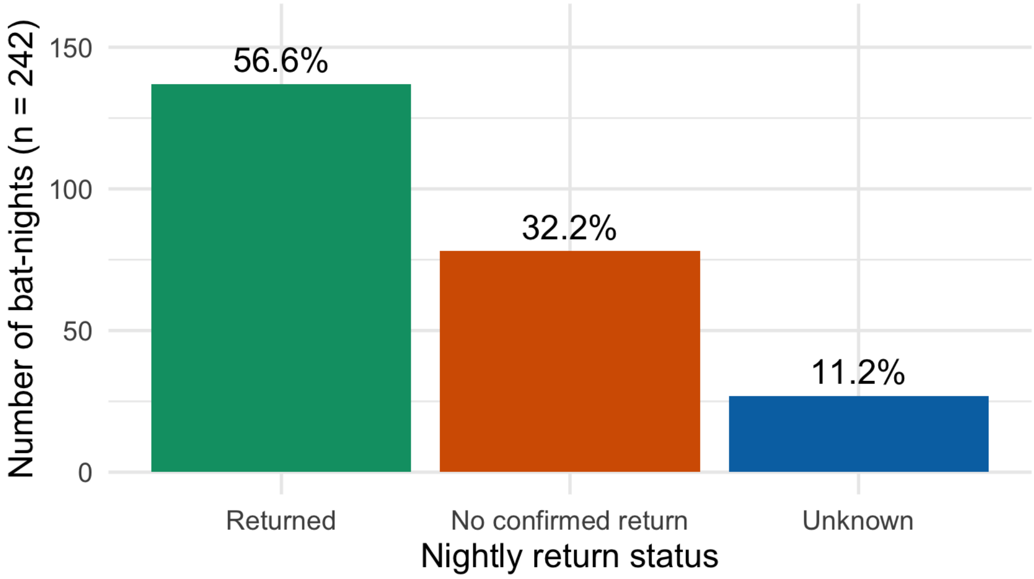

**Figure S2: Nightly return status across 242 bat-nights.** Bars show the number and percentage of bat-nights classified as 'Returned' (nights were a return to the focal roost could be confirmed from the final GPS fix or from the first GPS fix of the following night), 'No confirmed return' (nights were available data didn't confirm a return) or 'Unknown' (nights with insufficient data to determine return status, e.g., missing end-of-night or next-night fixes). Percentages are calculated over all 242 bat-nights. Excluding the 11% of bat-nights with unknown status, the return rate is 64% (137/215).

**Table S3: Model parameter priors and hyperpriors, and posteriors.**

| Parameter | Definition | Distribution | Prior | Posterior mean [95% credible intervals] | Effective sample size | psrf |
|---|---|---|---|---|---|---|
| $\sigma_1$ | Standard deviation for individual random effect | Gamma | Mean = 10 Variance = 10 | 2.67 [1.70, 3.67] | 32933 | 1.001 |
| $\theta_0$ | Intercept departure instance | Normal | Mean = 0 Variance = 1 | -2.40 [-3.49, -1.40] | 16040 | 1.001 |
| $\sigma$ | Standard deviation of departure instance | Beta | Mean = 0.09 Variance = 0.007 | 0.24 [0.15, 0.34] | 15431 | 1.005 |
| $\beta_0$ | Intercept in distance model | Normal | Mean = 2.3 Variance = 0.25 | 0.98 [0.79, 1.16] | 5008 | 1.001 |
| $\varphi$ | Overdispersion | Beta | Mean = $1^{-5}$ Variance = $1^{-9}$ | 0.258 [0.252, 0.264] | 16270 | 1.065 |
| $\rho$ | Probability of retention | Beta | Mean = 0.25 Variance = 0.01 | 0.29 [0.1, 0.49] | 34474 | 1.001 |
| $\tau_1$ | Precision for spike and slab prior | Gamma | Mean = 1000 Variance = 1,000,000 | 845.90 [2.56, 2392.23] | 28039 | 1.005 |
| $\tau_{ni}$ | Precision for coefficients of the interaction night and bat | Gamma | Mean = 1000 Variance = 1,000,000 | 1.24 [0.49, 2.10] | 5788 | 1.001 |

*Psrf* is the potential scale reduction factor. We extracted posterior means, effective sample sizes, and psrf estimates from the fitted model. Convergence was achieved for all parameters (psrf < 1.1).

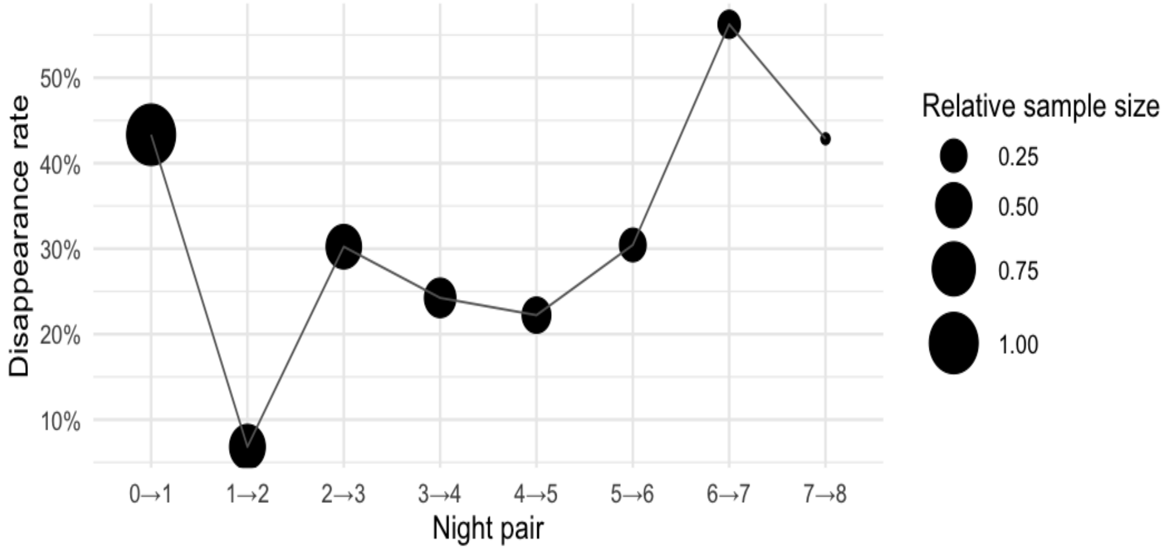


**Figure S3: Bat disappearance after the direct disturbance (nights 0-1) and bat detection loss across all the other nights.** Bat disappearance rates were highest between nights 0-1 and immediately decreased between nights 1-2. Higher rates on nights 6-7 and 7-8 were likely due to small sample sizes.

**Table S4: Results for the binomial generalised linear model on drivers of vampire bat (*Desmodus rotundus*) roost switching.**

Coefficients are on the log-odds scale. In a total of 93 vampire bats, 13 bats were excluded (three bats that died, one sessile bat/tag, and nine bats captured in roost 2, where we faced challenges of data recovery). A total of 80 bats were used in this analysis, where 31 bats didn't return to the roost (i.e., disappeared immediately after the direct human disturbance, with or without data for the night of disturbance), and 49 returned to the roost (i.e., had data for further nights). Table ordered by best-fitting model as determined by AIC.

| Model | Variable | Estimate | s.e. | P | AIC | PseudoR$^2$ |
|---|---|---|---|---|---|---|
| **M12** | SexM | -1.55 | 0.83 | 0.06 | 87.56 | 0.27 |
| | API155 | 2.97 | 0.88 | 0.0008 | | |

| | | | | | | |
|---|---|---|---|---|---|---|
| | AYA20 | 1.04 | 0.74 | 0.16 | | |
| | CUS6 | 2.65 | 0.87 | 0.002 | | |
| | Forearm | -1.10 | 0.42 | 0.008 | | |
| **M8** | API155 | 2.74 | 0.84 | 0.001 | 89.6 | 0.23 |
| | AYA20 | 0.92 | 0.71 | 0.20 | | |
| | CUS6 | 2.55 | 0.85 | 0.003 | | |
| | Forearm | -0.59 | 0.29 | 0.04 | | |
| **M13** | API155 | 2.77 | 0.85 | 0.001 | 91.17 | 0.24 |
| | AYA20 | 0.97 | 0.72 | 0.18 | | |
| | CUS6 | 2.48 | 0.85 | 0.004 | | |
| | Tag percentage | -0.19 | 0.30 | 0.52 | | |
| | Forearm | -0.63 | 0.30 | 0.035 | | |
| **M4** | API155 | 2.77 | 0.81 | 0.0007 | 91.98 | 0.19 |
| | AYA20 | 1.19 | 0.68 | 0.078 | | |
| | CUS6 | 2.64 | 0.82 | 0.0013 | | |
| **M9** | API155 | 2.80 | 0.82 | 0.0007 | 93.9 | 0.19 |
| | AYA20 | 1.23 | 0.69 | 0.075 | | |
| | CUS6 | 2.66 | 0.83 | 0.001 | | |
| | SexM | -0.15 | 0.55 | 0.79 | | |
| **M10** | API155 | 2.79 | 0.82 | 0.0006 | 93.9 | 0.19 |
| | AYA20 | 1.22 | 0.68 | 0.075 | | |
| | CUS6 | 2.62 | 0.82 | 0.002 | | |
| | Tag percentage | -0.07 | 0.28 | 0.80 | | |
| **M14** | API155 | 2.81 | 0.83 | 0.0007 | 95.87 | 0.19 |
| | AYA20 | 1.24 | 0.69 | 0.07 | | |
| | CUS6 | 2.64 | 0.83 | 0.0015 | | |
| | Tag percentage | -0.06 | 0.29 | 0.84 | | |
| | SexM | -0.12 | 0.56 | 0.82 | | |
| **M7** | SexM | -1.11 | 0.66 | 0.09 | 100.85 | 0.09 |
| | Forearm | -0.96 | 0.35 | 0.005 | | |
| **M3** | Forearm | -0.59 | 0.25 | 0.02 | 101.87 | 0.06 |
| **M11** | SexM | -1.05 | 0.67 | 0.12 | 102.31 | 0.09 |
| | Tag percentage | -0.19 | 0.26 | 0.47 | | |

| | | | | | | |
|---|---|---|---|---|---|---|
| | Forearm | -0.98 | 0.35 | 0.005 | | |
| **M5** | Forearm | -0.64 | 0.26 | 0.01 | 102.93 | 0.07 |
| | Tag percentage | -0.24 | 0.26 | 0.34 | | |
| **M2** | Tag percentage | -0.11 | 0.24 | 0.64 | 107.58 | 0.002 |
| **M1** | SexM | 0.11 | 0.47 | 0.81 | 107.75 | 5e-04 |
| **M6** | SexM | 0.16 | 0.48 | 0.73 | 109.46 | 0.003 |
| | Tag percentage | -0.13 | 0.24 | 0.60 | | |

Standard error (s.e.), P-value (P), Akaike Information Criterion (AIC), McFadden Pseudo$R^2$.

Tag percentage refers to the percentage of body mass that the tag represents. Kruskal-Wallis tests did not find significant differences in forearm length (Chi-squared = 4.84, df = 3, P = 0.18) neither in tag burden (Chi-squared = 3.00, df = 3, P = 0.39) between the four colonies. Sex ratios did not differ between roosts (Chi-squared = 2.21, df = 3, P = 0.53) and neither between returning and non-returning bats (Chi-squared = 9 e-06, df = 1, P = 0.998).

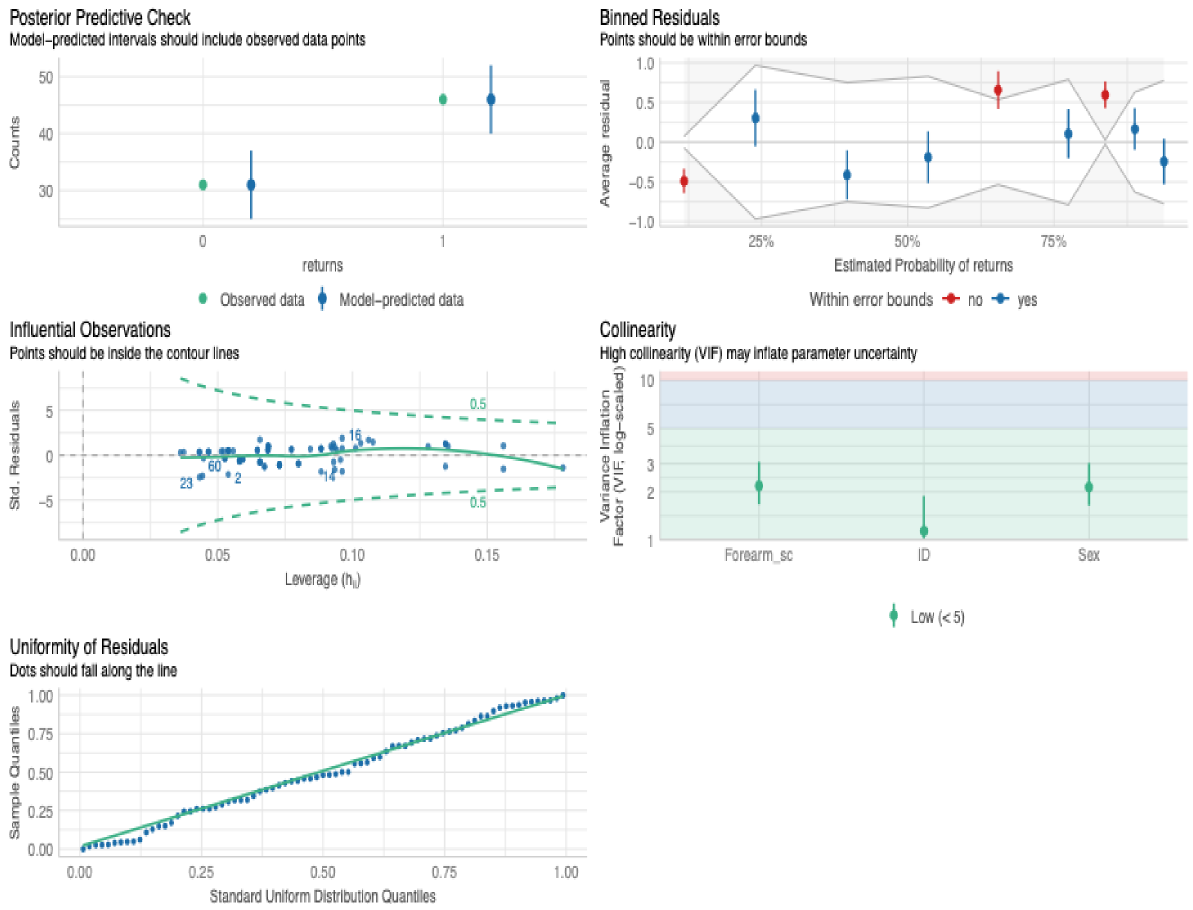


**Figure S4: Assessment of performance of binomial generalised linear model M12**. This plot used the R package performance (0.13.0) [8].

**Table S5: Results for the binomial generalised linear model on drivers of vampire bat (*Desmodus rotundus*) roost switching - interactions with bat traits.**

| Model | Variable | Estimate | s.e. | P | AIC | PseudoR$^2$ |
|---|---|---|---|---|---|---|
| **M2** | SexM | -1.07 | 0.66 | 0.11 | 101.86 | 0.1 |
| | Forearm | -0.64 | 0.45 | 0.16 | | |
| | SexM:Forearm | -0.69 | 0.70 | 0.32 | | |
| **M4** | SexM | -0.98 | 0.70 | 0.16 | 102.17 | 0.11 |
| | Tag percentage | 0.25 | 0.40 | 0.54 | | |
| | Forearm | -0.91 | 0.35 | 0.01 | | |
| | SexM:Tag percentage | -0.79 | 0.56 | 0.16 | | |
| **M5** | SexM | -1.03 | 0.67 | 0.12 | 103.50 | 0.1 |

| | | | | | | |
|---|---|---|---|---|---|---|
| | Forearm | -0.69 | 0.46 | 0.14 | | |
| | Tag percentage | -0.15 | 0.26 | 0.55 | | |
| | SexM:Forearm | -0.63 | 0.71 | 0.37 | | |
| **M6** | Tag percentage | -0.22 | 0.27 | 0.42 | 103.78 | 0.1 |
| | Forearm | -1.00 | 0.35 | 0.004 | | |
| | SexM | -1.11 | 0.68 | 0.10 | | |
| | Tag percentage:Forearm | 0.20 | 0.28 | 0.48 | | |
| **M3** | Tag percentage | -0.28 | 0.27 | 0.30 | 104.63 | 0.07 |
| | Forearm | -0.65 | 0.26 | 0.01 | | |
| | Tag percentage:Forearm | 0.15 | 0.27 | 0.59 | | |
| **M7** | SexM | -0.95 | 0.71 | 0.18 | 107.00 | 0.12 |
| | Tag percentage | 0.43 | 0.61 | 0.49 | | |
| | Forearm | -0.57 | 0.47 | 0.22 | | |
| | SexM:Tag percentage | -0.96 | 0.80 | 0.23 | | |
| | SexM:Forearm | -0.76 | 0.79 | 0.34 | | |
| | Tag percentage:Forearm | -0.14 | 0.50 | 0.78 | | |
| | SexM:Tag percentage:Forearm | 0.17 | 0.88 | 0.85 | | |
| **M1** | SexM | 0.19 | 0.51 | 0.71 | 107.68 | 0.04 |
| | Tag percentage | 0.40 | 0.38 | 0.29 | | |
| | SexM:Tag percentage | -1.01 | 0.55 | 0.06 | | |

Standard error (s.e.), P-value (P), Akaike Information Criterion (AIC), McFadden Pseudo$R^2$.

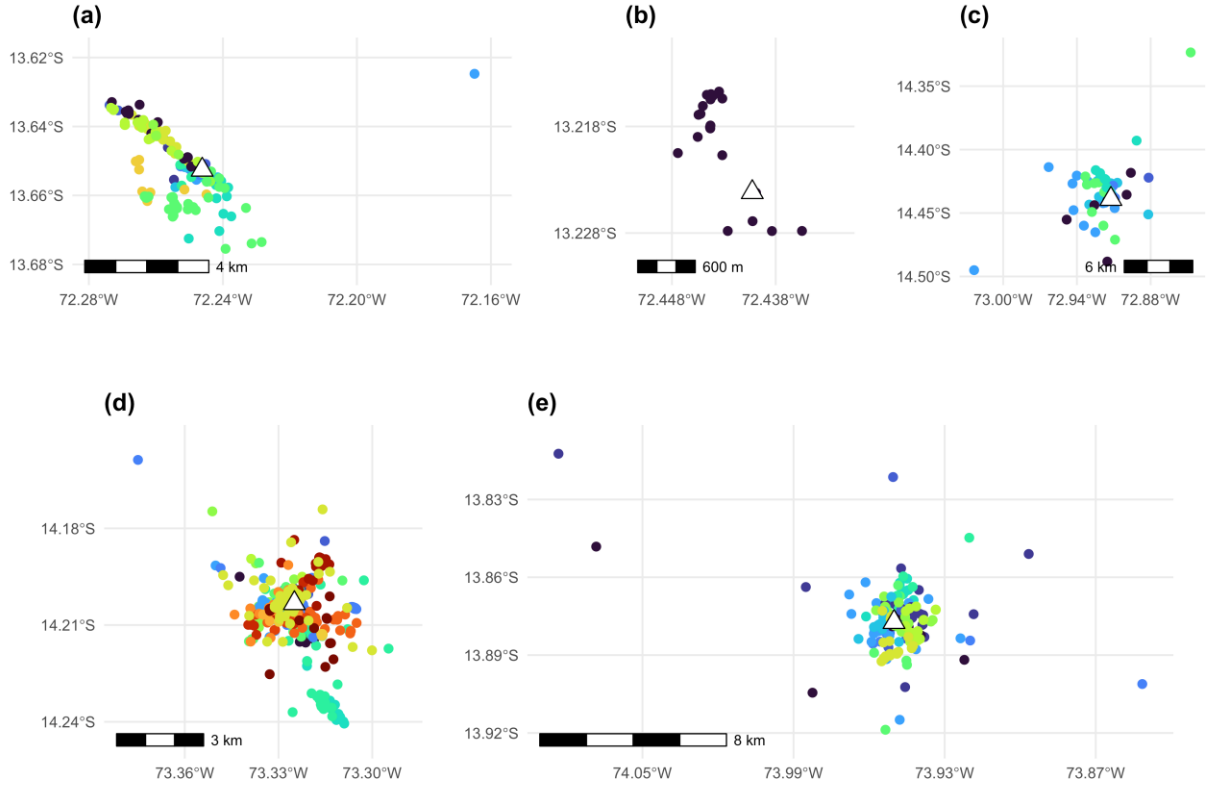


**Figure S5: Spatial distribution of vampire bat (*Desmodus rotundus*) GPS locations for five roosts in southern Peru.**

Data cleaning excluded a further 22.3% of the 1,700 fixes available (spatial and temporal filtering 18.7%; data quality attributes 5.8%, and unrealistic movements 75.5%), leaving 1,321 fixes from 55 bats. (a) roost 1; (b) roost 2; (c) roost 3; (d) roost 4; (e) roost 5. Each colour represents one individual bat. The white triangle represents the roost. Due to the varying scales of movement across the colonies, this figure does not use a standardised scale.

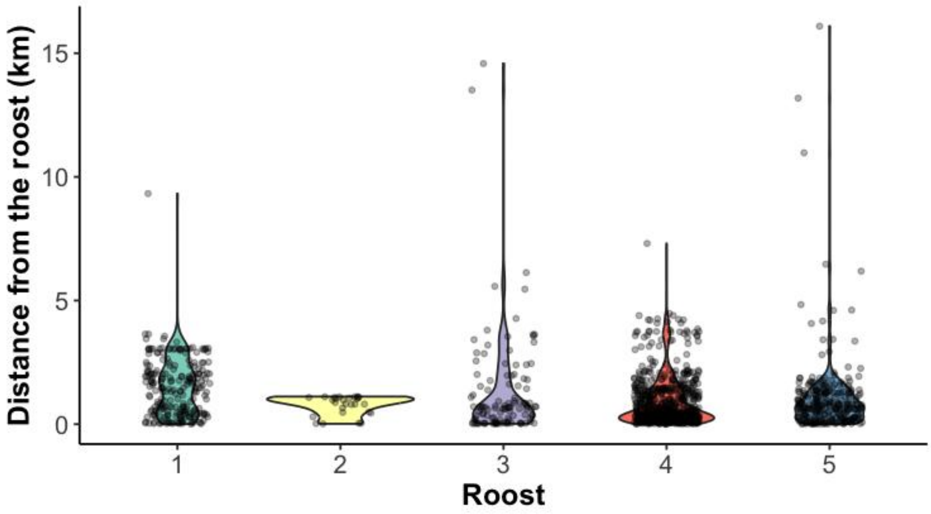


**Figure S6: Distribution of distance from the roost across roosts.**

Violin plots show the distribution of distance from the roost across roosts (one colour, one roost). The grey points are the vampire bat distances from the roost at each GPS fix.

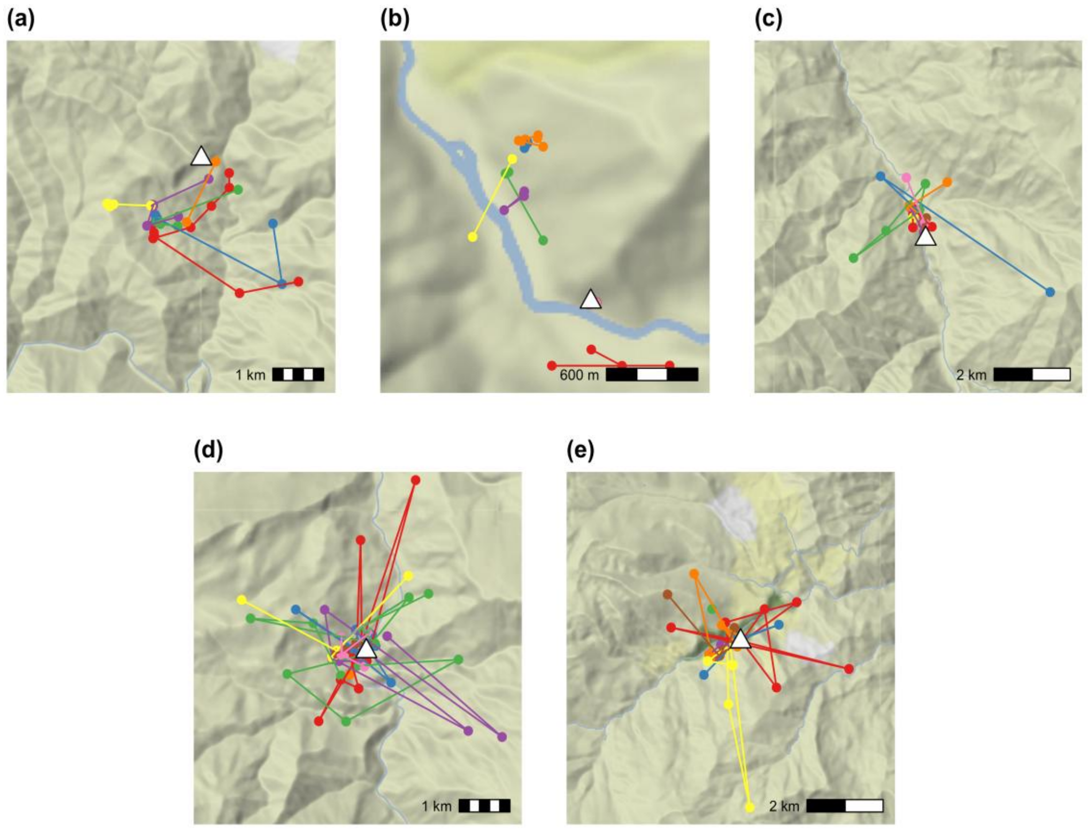


**Figure S7: Individual GPS data for vampire bats (*Desmodus rotundus*) for up to nine nights of tracking.**

One individual in each roost. (a) roost 1, (b) roost 2, (c) roost 3, (d) roost 4, and (e) roost 5. Each colour represents one night. The white triangle represents the roost. Due to the varying scales of movement across the colonies, this figure does not use a standardised scale. Maps used ggmap [9]. Base map tiles show terrain. Data © OpenStreetMap contributors, © Stadia Maps, © Stamen Design, © OpenMapTiles. Licensed under CC BY 3.0 / ODbL (https://www.openstreetmap.org/copyright).

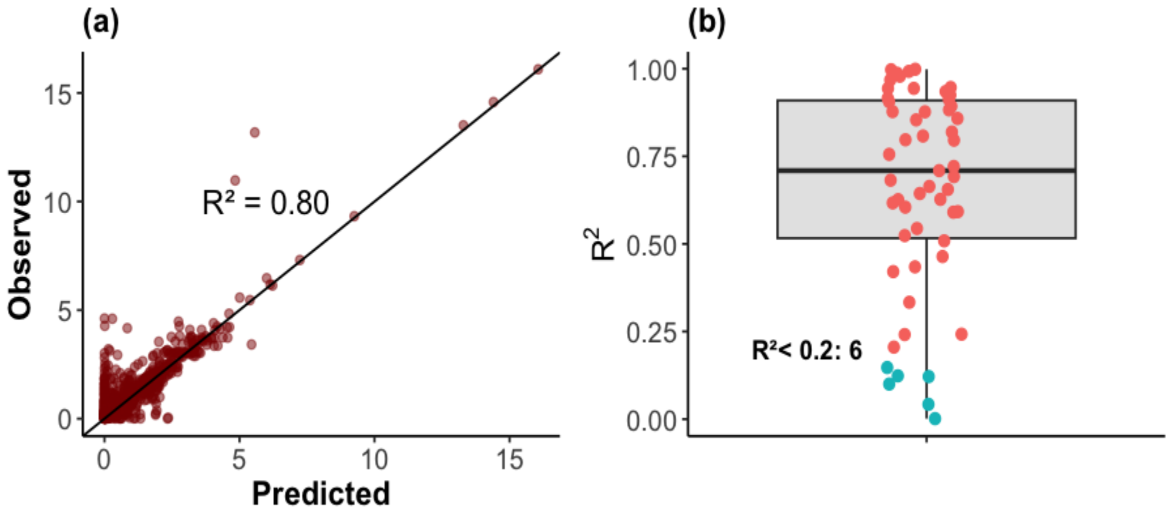


**Figure S8: Model goodness of fit for correlates of bat departure time and nightly distance from the roost.**

Results from the Bayesian state-space model. (a) Overall and (b) individual $R^2$. The model fitted well for most vampire bats, with a median individual $R^2$ of 0.71. Only six vampire bats (10.9%, out of 55) were poorly fit ($R^2$ <0.20). $R^2$ calculated using the R package metrica (2.1.0) [10], comparing observed distances with the posterior predicted mean.

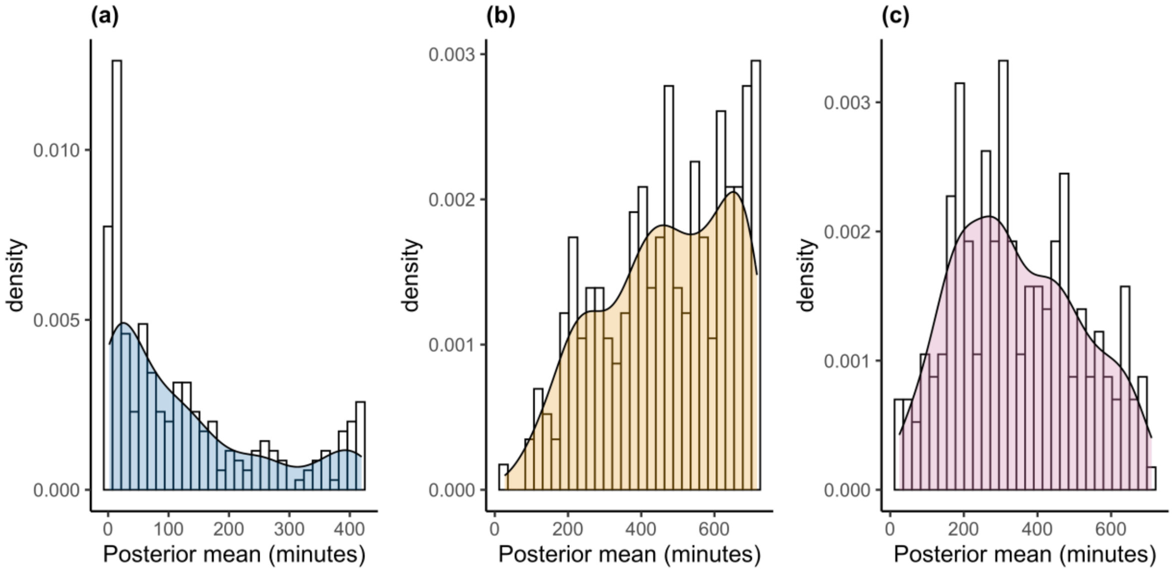


**Figure S9: Predicted times from the Bayesian state-space model.**

(a) Predicted departure time of vampire bats from the roost, (b) predicted return time, and (c) predicted duration of activity.

**Table S6: Summary of the Bayesian state-space model for vampire bats (*Desmodus rotundus*).**

Delta is the probability of a variable being kept in the model. The posterior mean, median, lower and upper bounds of the 95% credible interval, effective sample size (SSeff), autocorrelation at lag 2000 (AC.2000), and potential scale reduction factor (psrf) are displayed. All reported model parameters showed good convergence, with psrf values close to 1 (< 1.1).

| Model | Name | Delta | Mean | Median | Lower95 | Upper95 | SSeff | AC.2000 | psrf |
|---|---|---|---|---|---|---|---|---|---|
| Distance | Time post-disturbance (hours) | 0.95 | -0.037 | -0.039 | -0.069 | -0.008 | 20654 | 0.013 | 1.045 |
| Distance | Cattle density | 0.36 | -0.018 | -0.013 | -0.148 | 0.096 | 14870 | 0.003 | 1.002 |
| Distance | Forearm length | 0.31 | -0.010 | -0.008 | -0.130 | 0.098 | 20931 | 0.000 | 1.001 |
| Departure time | Sex | 0.31 | -0.010 | -0.007 | -0.132 | 0.108 | 38918 | 0.006 | 1.002 |
| Distance | Sex | 0.28 | -0.002 | -0.001 | -0.115 | 0.112 | 34856 | -0.003 | 1.000 |
| Distance | Roost density | 0.27 | 0.001 | 0.000 | -0.107 | 0.111 | 36280 | -0.003 | 1.000 |
| Distance | Proportion of permanent water | 0.27 | 0.003 | 0.003 | -0.103 | 0.117 | 29136 | 0.011 | 1.001 |
| Distance | Rural communities | 0.26 | -0.003 | -0.002 | -0.114 | 0.103 | 30466 | 0.007 | 1.001 |
| Departure time | Hours until dark | 0.19 | 0.009 | 0.006 | 0.000 | 0.028 | 39097 | 0.005 | 1.000 |

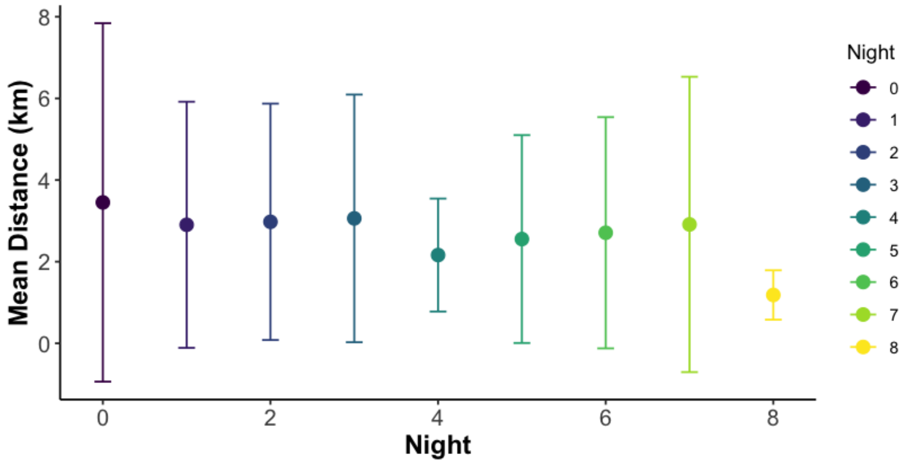


**Figure S10: Changes in the maximum predicted travel distance of vampire bats (*Desmodus rotundus*) over nights post-disturbance.**

Error bars represent one standard deviation (SD) around the mean of the maximum distance per night. Night 0 (mean = 3.45, SD = 4.39), night 1 (mean = 2.90, SD = 3.01), night 2 (mean = 2.98, SD = 2.90), night 3 (mean = 3.06, SD = 3.03), night 4 (mean = 2.16, SD = 1.38), night 5 (mean = 2.55, SD = 2.55), night 6 (mean = 2.71, SD = 2.83), night 7 (mean = 2.91, SD = 3.62), night 8 (mean = 1.18, SD = 0.61). The average decrease in maximum nightly distance over the 9 nights was 0.17 km (+- 0.06, P =0.035, $R^2$ = 0.49).

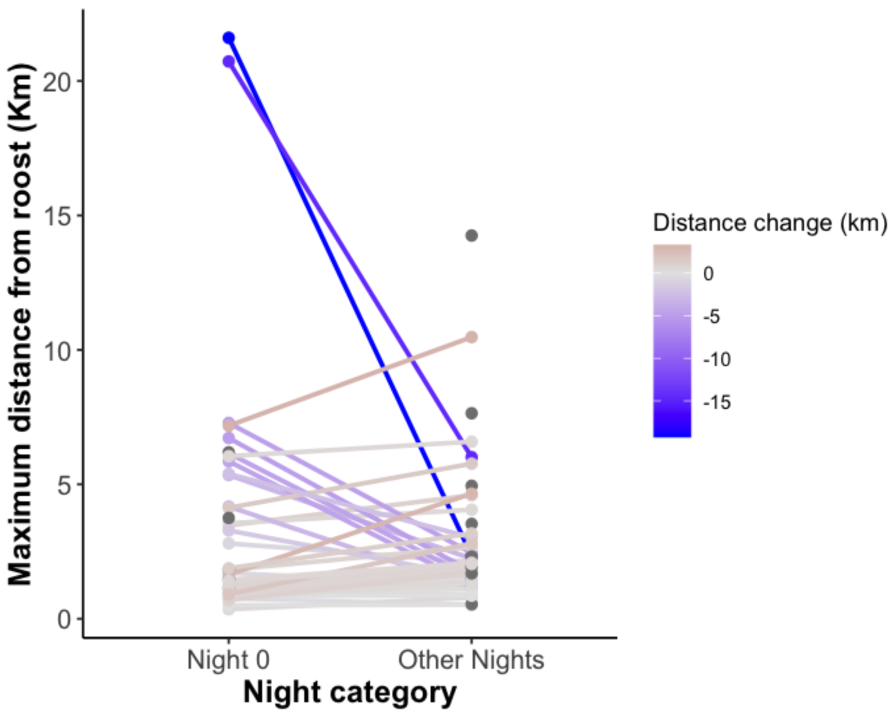


**Figure S11: Changes in the maximum predicted travel distance of vampire bats (*Desmodus rotundus*) from the roost**.

This plot compares 'night 0' with all subsequent night's post-disturbance. Each dot represents one bat. Bats that have only 'night 0' or no data for 'night 0' are coloured in dark grey.

**Table S7: Posterior summaries of the interaction coefficients between 55 vampire bats (*Desmodus rotundus*) and the night of disturbance.**

The posterior mean, median, lower and upper bounds of the 95% credible interval, effective sample size (SSeff), autocorrelation at lag 2000 (AC.2000), potential scale reduction factor (psrf), and direction of effect (positive or negative) are displayed. Name corresponds to each bat-ID. Convergence was achieved for all interaction coefficients (psrf < 1.1).

| Name | Mean | Median | Lower95 | Upper95 | SSeff | AC.2000 | psrf | type |
|---|---|---|---|---|---|---|---|---|
| 1 | 0.238 | 0.162 | -0.583 | 1.200 | 31,576 | -0.002 | 1.000 | Positive |
| 2 | -0.465 | -0.473 | -2.358 | 1.440 | 38,245 | 0.003 | 1.000 | Negative |
| 3 | 0.113 | 0.096 | -1.219 | 1.595 | 34,842 | -0.001 | 1.000 | Positive |
| 4 | -0.033 | -0.106 | -1.197 | 1.318 | 650 | 0.314 | 1.011 | Negative |
| 5 | -0.309 | -0.309 | -0.410 | -0.214 | 40,000 | 0.001 | 1.002 | Negative |
| 6 | 1.032 | 1.007 | 0.744 | 1.393 | 39,499 | 0.001 | 1.002 | Positive |
| 7 | 0.473 | 0.437 | -0.849 | 1.900 | 36,730 | 0.004 | 1.000 | Positive |
| 8 | -0.203 | -0.205 | -0.424 | 0.022 | 39,874 | -0.007 | 1.000 | Negative |
| 9 | -0.459 | -0.466 | -2.256 | 1.331 | 39,691 | 0.004 | 1.000 | Negative |
| 10 | -0.008 | -0.058 | -0.372 | 0.501 | 39,678 | -0.004 | 1.000 | Negative |
| 11 | 2.151 | 2.099 | 1.654 | 2.774 | 14,865 | 0.019 | 1.001 | Positive |
| 12 | 1.457 | 1.457 | 1.250 | 1.657 | 39,693 | -0.004 | 1.001 | Positive |
| 13 | -0.098 | -0.092 | -2.004 | 1.768 | 39,213 | -0.007 | 1.000 | Negative |
| 14 | 0.953 | 0.928 | -0.239 | 2.192 | 9,860 | 0.066 | 1.008 | Positive |
| 15 | -0.353 | -0.333 | -0.804 | 0.073 | 30,021 | 0.134 | 1.048 | Negative |
| 16 | 1.339 | 1.188 | 0.713 | 2.403 | 388 | 0.782 | 1.004 | Positive |
| 17 | -0.001 | 0.000 | -1.897 | 1.895 | 40,658 | 0.003 | 1.000 | Negative |
| 18 | 0.719 | 0.723 | -0.308 | 1.784 | 10,435 | 0.003 | 1.001 | Positive |
| 19 | 0.791 | 0.780 | -0.323 | 1.989 | 22,723 | -0.007 | 1.000 | Positive |
| 20 | -0.329 | -0.367 | -2.103 | 1.556 | 40,459 | -0.002 | 1.000 | Negative |
| 21 | -0.089 | -0.084 | -2.016 | 1.788 | 42,654 | 0.013 | 1.000 | Negative |
| 22 | -0.003 | 0.000 | -1.909 | 1.891 | 40,499 | 0.012 | 1.000 | Negative |
| 23 | -0.070 | -0.066 | -2.043 | 1.827 | 40,327 | -0.003 | 1.000 | Negative |
| 24 | 0.007 | 0.008 | -1.859 | 1.977 | 40,000 | 0.001 | 1.000 | Positive |
| 25 | -0.201 | -0.186 | -2.192 | 1.745 | 39,937 | 0.000 | 1.000 | Negative |
| 26 | -0.530 | -0.611 | -2.102 | 1.258 | 39,731 | -0.011 | 1.000 | Negative |
| 27 | 0.004 | 0.001 | -1.877 | 1.925 | 39,764 | 0.005 | 1.000 | Positive |
| 28 | 0.346 | 0.338 | -0.742 | 1.482 | 35,335 | -0.001 | 1.000 | Positive |
| 29 | -0.105 | -0.103 | -2.040 | 1.739 | 40,000 | -0.003 | 1.000 | Negative |
| 30 | -0.004 | -0.007 | -1.882 | 1.909 | 39,640 | -0.002 | 1.000 | Negative |
| 31 | -0.308 | -0.332 | -2.099 | 1.632 | 26,312 | 0.011 | 1.000 | Negative |

| Name | Mean | Median | Lower95 | Upper95 | SSeff | AC.2000 | psrf | type |
|---|---|---|---|---|---|---|---|---|
| 32 | 0.294 | 0.360 | -0.557 | 0.965 | 21,548 | -0.003 | 1.001 | Positive |
| 33 | -0.559 | -0.561 | -0.780 | -0.352 | 6,925 | 0.094 | 1.006 | Negative |
| 34 | -0.601 | -0.547 | -1.419 | 0.075 | 33,323 | 0.002 | 1.000 | Negative |
| 35 | -0.216 | -0.217 | -2.215 | 1.734 | 37,504 | -0.005 | 1.000 | Negative |
| 36 | -0.401 | -0.469 | -1.776 | 1.261 | 37,245 | 0.008 | 1.000 | Negative |
| 37 | -0.259 | -0.260 | -0.526 | 0.011 | 39,655 | -0.004 | 1.001 | Negative |
| 38 | -0.528 | -0.523 | -1.564 | 0.517 | 31,131 | 0.016 | 1.000 | Negative |
| 39 | -0.025 | -0.039 | -1.610 | 1.523 | 38,021 | 0.000 | 1.000 | Negative |
| 40 | -1.318 | -1.309 | -1.802 | -0.858 | 30,643 | 0.006 | 1.002 | Negative |
| 41 | 0.686 | 0.657 | -0.073 | 1.507 | 3,738 | 0.151 | 1.000 | Positive |
| 42 | -0.500 | -0.498 | -1.566 | 0.621 | 35,612 | -0.007 | 1.000 | Negative |
| 43 | 0.000 | -0.001 | -1.866 | 1.927 | 39,224 | 0.001 | 1.000 | Negative |
| 44 | -0.962 | -0.979 | -2.312 | 0.381 | 4,232 | 0.099 | 1.012 | Negative |
| 45 | -0.140 | -0.133 | -2.099 | 1.744 | 34,560 | -0.006 | 1.000 | Negative |
| 46 | -0.130 | -0.112 | -2.120 | 1.726 | 40,000 | 0.010 | 1.000 | Negative |
| 47 | -0.004 | -0.013 | -1.938 | 1.882 | 40,000 | 0.008 | 1.000 | Negative |
| 48 | -0.073 | -0.075 | -1.964 | 1.808 | 39,705 | 0.007 | 1.000 | Negative |
| 49 | 0.004 | 0.004 | -1.838 | 1.918 | 40,561 | 0.010 | 1.000 | Positive |
| 50 | -0.083 | -0.076 | -1.985 | 1.879 | 39,435 | -0.003 | 1.000 | Negative |
| 51 | -0.001 | 0.001 | -1.948 | 1.868 | 40,000 | 0.004 | 1.000 | Negative |
| 52 | -0.390 | -0.411 | -2.208 | 1.523 | 39,595 | -0.012 | 1.000 | Negative |
| 53 | 0.007 | 0.013 | -1.856 | 1.919 | 40,000 | 0.000 | 1.000 | Positive |
| 54 | 0.969 | 0.912 | -0.269 | 2.335 | 4,955 | 0.130 | 1.000 | Positive |
| 55 | -0.143 | -0.149 | -2.009 | 1.673 | 39,562 | 0.004 | 1.000 | Negative |

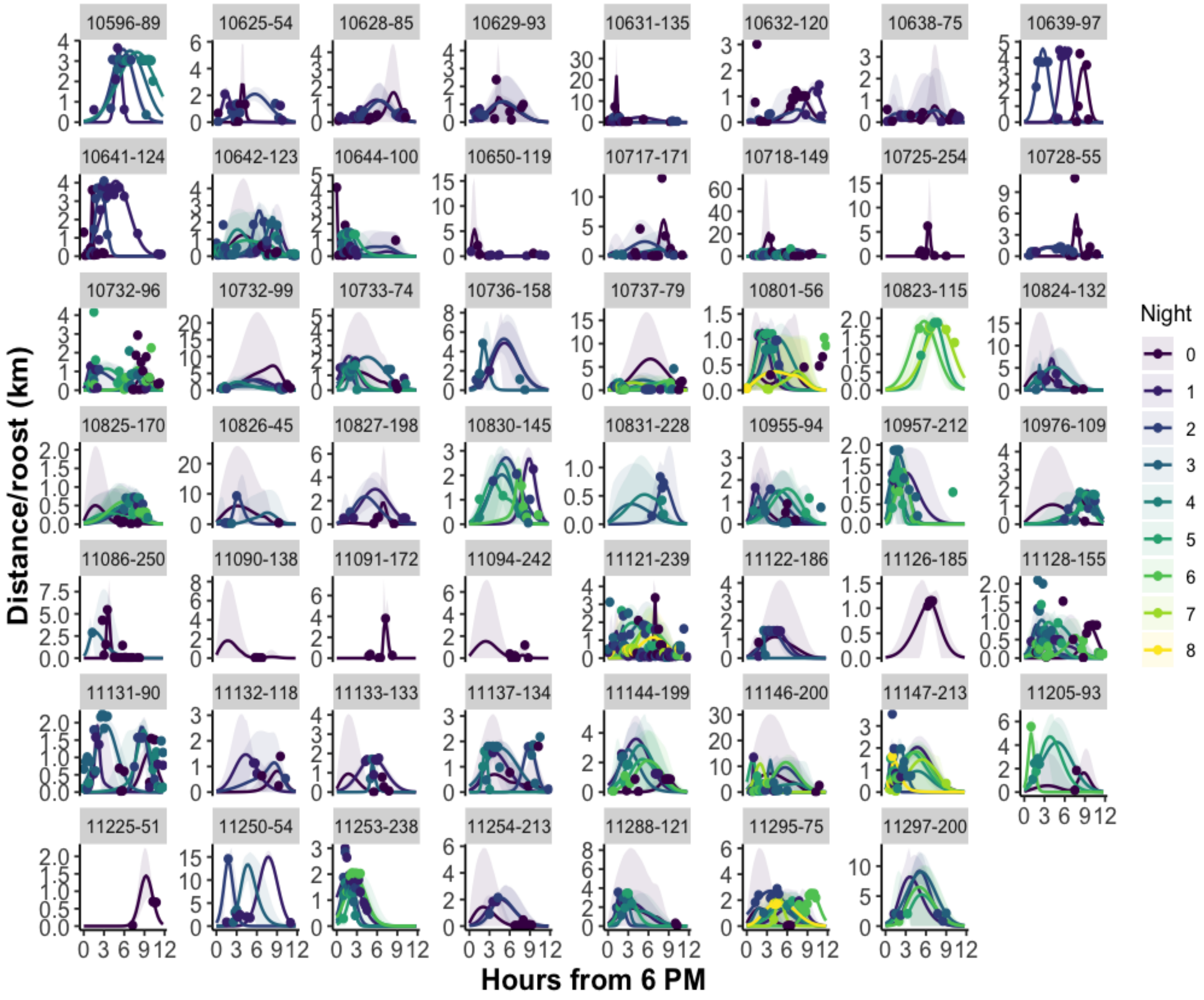


**Figure S12: Reconstructed distance-over-time curves for 55 vampire bats (*Desmodus rotundus*).** Individual and night variation. Predicted distances from the roost (posterior mean, y-axis) and time since 6 PM (x-axis). Each colour represents one night. The shaded area indicates the credible intervals, and the dots the observed values. Due to the varying scales of movement across the colonies, the y-axis is not standardised. In this figure, we would like to point out the model appearance of bimodal curves (two trips) being performed in 13% of the bat-nights.

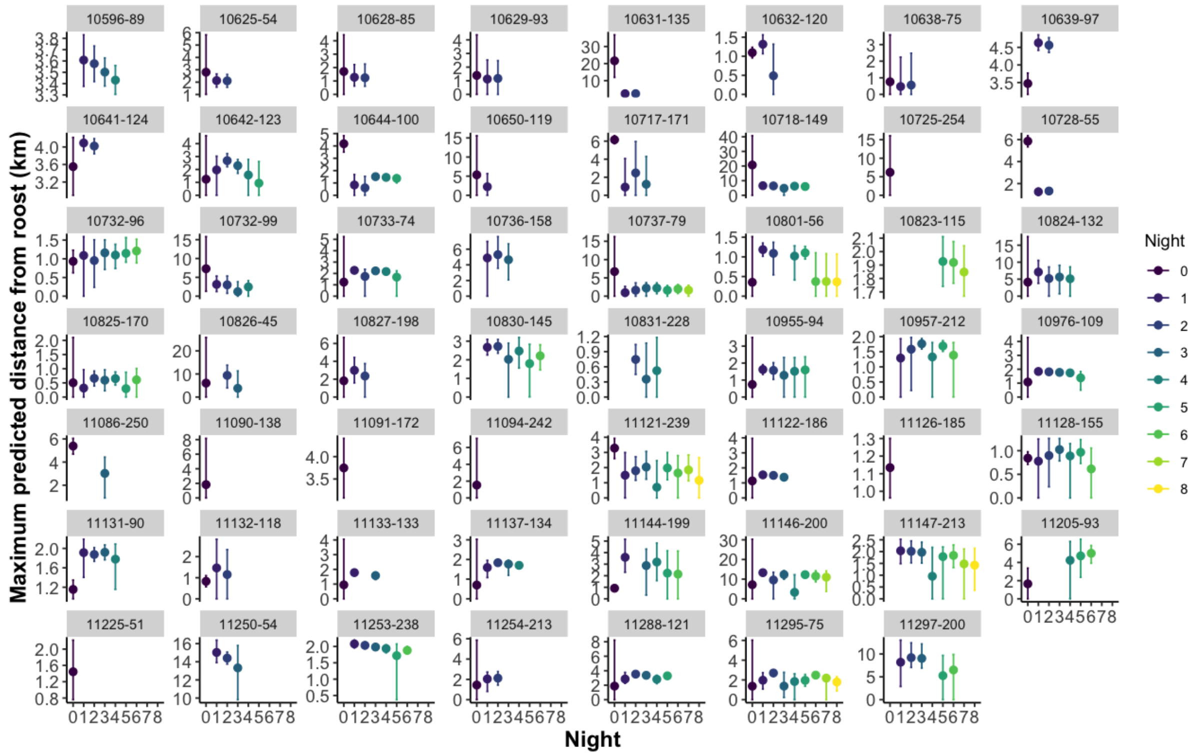


**Figure S13: Nightly individual distance effects.**

Maximum predicted distances (with credible intervals) for each individual and night. The y-axis represents the maximum predicted distances from the roost (posterior mean), and the x-axis represents the night. Due to the varying scales of movement across the colonies, the y-axis is not standardised.

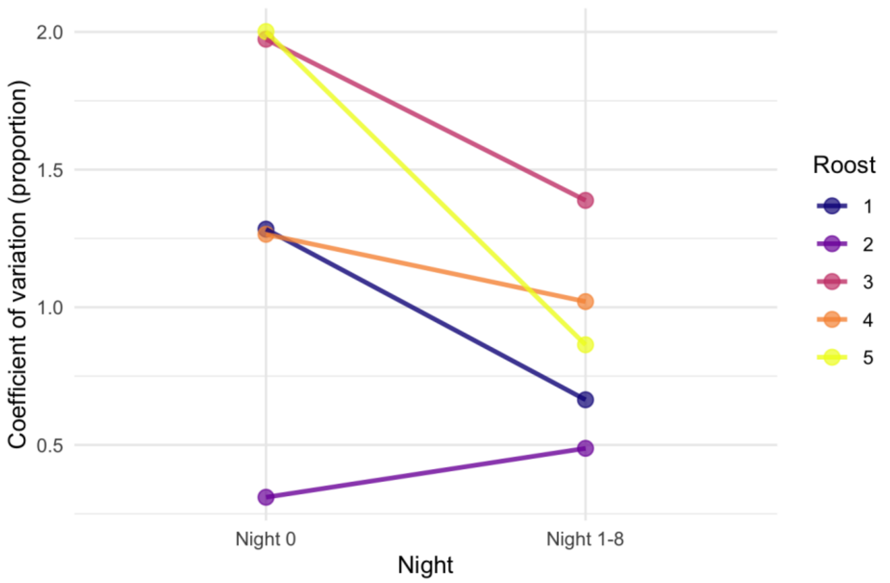


**Figure S14: Coefficient of variation of observed distance of vampire bats (*Desmodus rotundus*) from the roost by roost.**

Plot comparing 'night 0' with all the other nights (nights 1-8) for five roosts in southern Peru. Except for roost 2 (only one individual captured in roost 2, in one capture session), the variability in the distance from the roost was higher on 'night 0' than in all the other nights.
